# Dissociating Attentional Capture from Action Cancellation in the Stop Signal Task

**DOI:** 10.1101/2022.12.20.521300

**Authors:** Simon Weber, Sauro Salomoni, Callum Kilpatrick, Mark R. Hinder

**Affiliations:** Sensory Neuroscience and Aging Research Lab, the University of Tasmania, Churchill Ave, Hobart, Tasmania, Australia, 7005

**Keywords:** Response inhibition, aging, stop signal task, action cancellation, electromyography, stopping interference effect

## Abstract

Inhibiting ongoing responses when environmental demands change is a critical component of human motor control. Experimentally, the stop signal task (SST) represents the gold standard response inhibition paradigm. However, an emerging body of evidence suggests that the SST conflates two dissociable sources of inhibition, namely an involuntarily pause associated with attentional capture and the (subsequent) voluntary cancellation of action. The extent to which these processes also occur in other response tasks is unknown.

24 younger (20-35 years) and 23 older (60-85 years) adults completed a series of tasks involving rapid unimanual or bimanual responses to a visual stimulus. A subset of trials required cancellation of one component of an initial bimanual response (i.e., selective stop task; stop left response, continue with right response) or enacting an additional response (e.g., press left button as well as right button). Critically, both tasks involved some infrequent stimuli which bore no behavioural imperative (i.e., they had to be ignored).

EMG recordings of voluntary responses during the stopping tasks revealed bimanual covert responses (i.e., muscle activation which was suppressed before a button press ensued), consistent with a pause process, following both stop *and* ignore stimuli, before the required response was subsequently enacted. Critically, we also observed the behavioural consequences of a similar involuntary pause in trials where action cancellation was not part of the response set (i.e., when the additional stimulus required additional action or ignoring, but not inhibition). The findings shed new light on the mechanisms of inhibition and their generalisability to other task contexts.

## 1. Introduction

To safely navigate our environment, we often need to inhibit planned or ongoing actions in response to updated sensory input. The resulting inhibition may be ‘global’, resulting in the cancellation of all actions, or ‘selective’, whereby only a subset of our current movements is cancelled (Aron & Verbruggen, 2008). Many everyday scenarios require the latter kind of selective inhibition. For example, suppose you are driving around a bend when a car suddenly pulls onto the road up ahead. In that moment you must cancel one movement you were about to make (pressing down on the accelerator) while continuing another (shifting your hand to turn the wheel, to safely navigate the corner). Experimental evidence suggests that cancelling only a subset of ongoing actions, referred to as ‘motor selective stopping’ is performed sub-optimally: there is a small time cost afforded to any non-cancelled actions, whereby their execution is delayed (Aron & Verbruggen, 2008). The reasons for this phenomenon, and the contexts that give rise to it, are a current topic of investigation (For review, see Wadsley et al., 2022). Investigating these fundamental aspects of motor control across the lifespan informs our understanding of the physiological changes that take place with aging. Indeed, inhibitory control degrades as we age, conceivably impacting our capacity to safely and effectively interact with our complex world (Buchman et al., 2009; Coxon et al., 2016). However, the underlying mechanisms of selective stopping, and how these change in the later stages of adulthood, remain incompletely understood.

The Stop Signal Task (SST) is a behavioural paradigm which allows for characterisation of response cancellation processes (Logan & Cowan, 1984). While SSTs are largely heterogenous, they typically require the participant to rapidly enact a prescribed movement (e.g., press a button) in response to a visual cue. On a minority of trials this cue is followed by a stop signal (usually a visual or auditory cue), which indicates that the participant must cancel their initiated response (Verbruggen et al., 2019). In order to broaden our understanding of the processes underpinning inhibitory control, a number of variations of the standard SST have been developed. Of particular relevance are ‘motor selective’ SSTs, featuring stop trials in which the participant is required to only inhibit part of a multi-component ‘go’ response, while continuing with another component, e.g., stopping only one hand during a bimanual movement; (Aron & Verbruggen, 2008; Coxon et al., 2007, Wadsley et al., 2022). This experimental paradigm can be used to investigate the process of ‘motor selective stopping’ described above.

### 1.2. The Stopping Interference Effect

Research employing motor selective SSTs has consistently demonstrated that the reaction time (RT) of the continued action component (e.g., the RT of the hand that was *not* required to stop in successful selective stop trials) is slower (by ∼100ms, though results vary) than the RT observed in trials where no stop signal was presented (Cai et al., 2011; Claffey et al., 2010; Coxon et al., 2007; Drummond et al., 2018; Majid et al., 2012; Raud & Huster, 2017). This delay has been termed the ‘stopping interference’ (SI) effect (Aron & Verbruggen, 2008). The source of the SI effect is multifaceted and context sensitive. Potential mediators include delays associated with the decoupling of a functionally coupled motor program (Wadsley et al., 2019), and the presence of a global stop mechanism preceding the initiation of a new unimanual response (MacDonald et al., 2014, 2017). The SI effect may also be mediated by a transient global ‘pause’ which is distinct from the conscious cancellation of a movement, and could be associated with attentional capture to any unexpected and behaviourally relevant sensory stimuli (Diesburg & Wessel, 2021).

### 1.3. SSTs conflate two distinct sources of inhibition

A growing body of evidence suggests that unexpected events trigger transient and non-selective inhibition of motor output (Wessel & Aron, 2017). Cortical and subcortical regions associated with inhibition are recruited in response to infrequent or unexpected behavioural cues regardless of whether or not the behavioural imperative associated with such cues involves action cancellation (González-Villar et al., 2022; Sebastian et al., 2021; Tatz et al., 2021). These findings provide support for a recent pause-then-cancel model of action inhibition, which distinguishes between two complimentary inhibitory processes (Diesburg & Wessel, 2021; Schmidt & Berke, 2017). A pause process, mediated by the hyperdirect cortico-basal-ganglia pathway (Chen et al., 2020) generates a transient global suppression of motor activity that relates to attentional orienting. Following this, a distinct cancel process, generated via the slower indirect pathway (Aron, 2011), allows the selective action cancellation. Because the stop signals in an SST occur on a minority of trials and follow a variable delay, they will not only lead to voluntary action cancellation, but will initially also trigger attentional orienting (and the associated pause response). As a result, the SST conflates these two distinct sources of inhibition.

In an attempt to disentangle the aspects of the SI effect which are associated with the presentation of an additional stimulus, Ko & Miller (2013) combined motor selective stopping (described above) with ‘stimulus selective’ stopping (Bissett & Logan, 2014). Stimulus selective SSTs involve two perceptually distinct types of infrequent cue, only one of which is associated with the imperative to stop (van de Laar et al., 2011). The other cue (variably referred to as an ‘ignore cue’, a ‘continue cue’, or a ‘controlled go’) bears no behavioural imperative; the participant must simply enact their response as though no additional cue has appeared. Ko and Miller (2013) compared RTs in go trials (where only the ‘go’ cue appeared) to ‘controlled go’ trials (where an ‘ignore’ cue appeared) and observed a small, non-significant degree of slowing in the controlled go trials. They also observed that RTs in the continuing hand during successful selective stop trials were significantly longer than reactions in both go and controlled go trials. Based on these findings, the authors suggest that the majority of the SI effect is generated from mechanisms associated directly with the act of stopping, rather than attentional capture associated with the cues themselves.

Notably, however, Ko and Miller used a high rate of ‘controlled go’ and stop signals (stop signals occurred on 40% of trials, ignore cues occurred on 48% of trials and only 12% of trials did not have some kind of additional cue associated with them). Since the publication of this research there have been a number of findings which suggest that this high rate of additional signals could have attenuated the delays following presentation of an ignore stimulus. Specifically, the high rate of stop and ignore signals would increase participants’ expectation of a) the presentation of a second cue, and b) the requirement to stop (Verbruggen et al., 2019). The high proportion of stop and ignore stimuli may therefore have diminished the pause effect (which is more likely to occur in response to infrequent and unexpected stimuli; Diesburg & Wessel, 2021) and thus attenuated the SI effects.

The current experiment was designed to combine stimulus and motor selective stopping, but using infrequent (unexpected) stop and ignore signals (c.f. Ko & Miller, 2013). We anticipated a greater pause/slowing effect from ignore cues would be observed, suggesting that previous papers had underestimated this. We also sought to test whether an involuntary pause effect (previously considered as an aspect of action cancellation) could be similarly observed in a non-stopping context. That is, if a pause process (associated with attentional capture) contributes to the observed SI effects during a selective stop then a slowing effect similar to the SI effect may also be observable outside of a stopping context (e.g., in response to an additional signal presenting after the initial Go stimulus but which is associated with the requirement to execute an additional response rather than cancel a response). Given that a behavioural pause would have no benefit at all (in fact, a decrement) in movement tasks where stopping was not a possibility, this would provide a test of whether this mechanism is subconscious and not modifiable, and at play in a broader array of behavioural contexts where actions need to be updated, but not cancelled. The current experiment explores this possibility by comparing performance in tasks requiring unexpected stopping and unexpected additional actions, both of which involved ignore stimuli.

### 1.4. Characterising action cancellation using electromyography (EMG)

Conventional measures of reactive inhibition involve deriving an estimate of stop signal reaction time (SSRT) based on behavioural results. Many of these processes have recently been called into question based on violations of assumptions in the traditional horse race model underlying them (Gulberti et al., 2014; Verbruggen & Logan, 2015), especially in selective stopping contexts (Bissett et al., 2021). Furthermore, conventional SSRT calculations fail to compensate for inconsistencies in the attentional components of the stopping process (Matzke et al., 2017).

Directly recording muscle activity using electromyography (EMG) during an SST allows for precise single trial measures of inhibition latency and response amplitude (Raud & Huster, 2017). By analysing EMG recordings, the delay between the appearance of the stop signal and the time at which muscle activity of a partially initiated response begins to decrease can be directly calculated (Jana et al., 2020; Raud et al., 2022). This not only circumvents many of the limitations associated with traditional methods of estimating stopping latency, but also removes the ballistic stage of action cancellation (i.e., mechanical delays associated with muscle contraction to the point of a button press being registered), which is subject to variations in experimental conditions, such as button stiffness, the effector with which the participants execute the movement, and whether the response requires a button press or release (e.g. Coxon et al. 2007). Furthermore, SSRT estimates result in only a single value per participant, belying the fact that an individual’s stopping latency is likely to vary between trials. In contrast, EMG indices of inhibition latency represent a distribution of that individual’s stopping latency (Raud et al., 2022).

The current experiment extends recent work aiming to elucidate the processes underlying selective stopping by combining novel behavioural paradigms with single trial analysis of covert and overt muscle responses in young and older adults. We report that previous research underestimates the proportion of inhibition during action cancellation that is associated with attentional capture and that the notion of pausing in response to unexpected stimuli (conceivably to facilitate subsequent inhibition) also occurs when inhibition is *not* required.

## 2. Methods

### 2.1. Participants

52 volunteers took part in the study, though the data from five participants were excluded prior to analysis (three due to excessive noise in their EMG recordings and two due to significant behavioural deviations from task instructions). Following exclusions, the younger cohort consisted of 24 participants (mean age = 24.5 years, SD = 5.5, range = 18 – 27; 12 female, 12 male), and the older cohort consisted of 23 participants (mean age = 70.3 years, SD = 5.8, range = 61 – 83; 14 female, 9 male). The younger cohort was recruited via the University of Tasmania’s psychology research participation system, while the older participants were recruited via a group email sent to the University of the Third Age (U3A) in Hobart (Tasmania, Australia). As compensation for their time, participants received either two hours of course credit (for those requiring credit) or a $20 shopping voucher. All participants provided informed consent and had normal or corrected to normal vision. This research was approved by the Tasmania Human Research Ethics Committee #H001698.

### 2.2. Procedure

#### 2.2.1. Computer Interfaces

Participants were seated approximately 80cm from a computer monitor with their forearms pronated, with forearms and hands resting on a desk shoulder width apart; each index finger rested against one of two custom made response buttons. Button presses were registered via The Black Box Toolkit USB response pad and recorded using PsychoPy3 (Peirce et al., 2019). The buttons were mounted in the vertical plane, such that registering a response required participants to abduct their index fingers (inwards) such that the first dorsal interossei (FDI) muscle acted as an agonist to facilitate recordings of task related electromyographic activity. During the experiment, participants were required to press the button/s as quickly as possible in response to an initial imperative cue. In different conditions (described below) additional cues required cancellation of some part of the initial response, required additional responses, or simply had to be ignored). The total duration of the experiment was approximately two hours.

#### 2.2.2 Electromyography (EMG)

EMG recordings were made using disposable adhesive electrodes placed on the first dorsal interossei (FDI) of participants’ left and right index fingers. On each hand, two electrodes were placed in a belly-tendon montage, with a third ground electrode placed on the head of the right ulna. EMG data were recorded using the computer software Signal (Cambridge Electronic Design Ltd.). The analogue EMG signals were band-pass filtered at 20-1000 Hz, amplified 1000 times, and sampled at 2000Hz (see section 2.5 for details regarding EMG data processing). Participants were requested to remain as relaxed as possible between trials, and only activate muscles to press the buttons. Once the experiment commenced, the researcher would remind the participant to relax their hands between trials, if recordings became noisy with background activity.

### 2.3. Behavioural Tasks

#### 2.3.1. Overview

The experiment consisted of four conditions, designed to tease apart the contributions of attentional capture and action inhibition processes in selective stopping. To this end, we used tasks requiring both selective inhibition as well as a novel paradigm where instead of a stop signal appearing on infrequent trials, the unexpected stimulus required an additional action. These two variants were completed with, and without, additional ‘ignore’ stimuli, in counterbalanced order. Accordingly, the four conditions, explained in detail below, were **1**) stop signal task (SST); **2**) stop signal task with ‘ignore’ trials (SSTignore); **3**) ‘additional go task’ (AGT); and **4**) ‘additional go task’ with ‘ignore’ trials (AGTignore).

#### 2.3.2. Instructions, go only blocks and randomisation

Each of the four conditions started with a series of instruction slides appearing on the screen. Following this, participants completed a short block consisting only of ‘go’ trials (described below). Go trials in the SST and SSTignore condition required bimanual responses while those in the AGT and AGTignore required unimanual responses and occurred on either the left or the right. These trials served to provide a baseline measure of reaction time (RT) in the absence of any other task demands. During these blocks, trials which were very fast (RT < 100ms), or slow (RT > 450ms) were repeated, and participants continued until 10 appropriate trials were completed. In the SST and SSTignore conditions trials in which an asynchronous response was made (difference of > 50ms between responses) were also repeated.

All trials, irrespective of condition or trial type, began with a fixation cross appearing for a duration drawn from a truncated exponential distribution ranging from 250-1000ms. This served to reduce the predictability of the go-signal onset and ensure that participants were responding to the visual stimulus, rather than predicting its appearance (Verbruggen et al., 2019). The visual cues (see figure 1) were circles which were 3cm in diameter and 15 cm apart. The response window lasted 1200ms before a feedback screen displaying the RT of both the left and right hands appeared for 1000ms. Following this, the screen was blank for 200ms before the next trial began.

**Figure 1:**
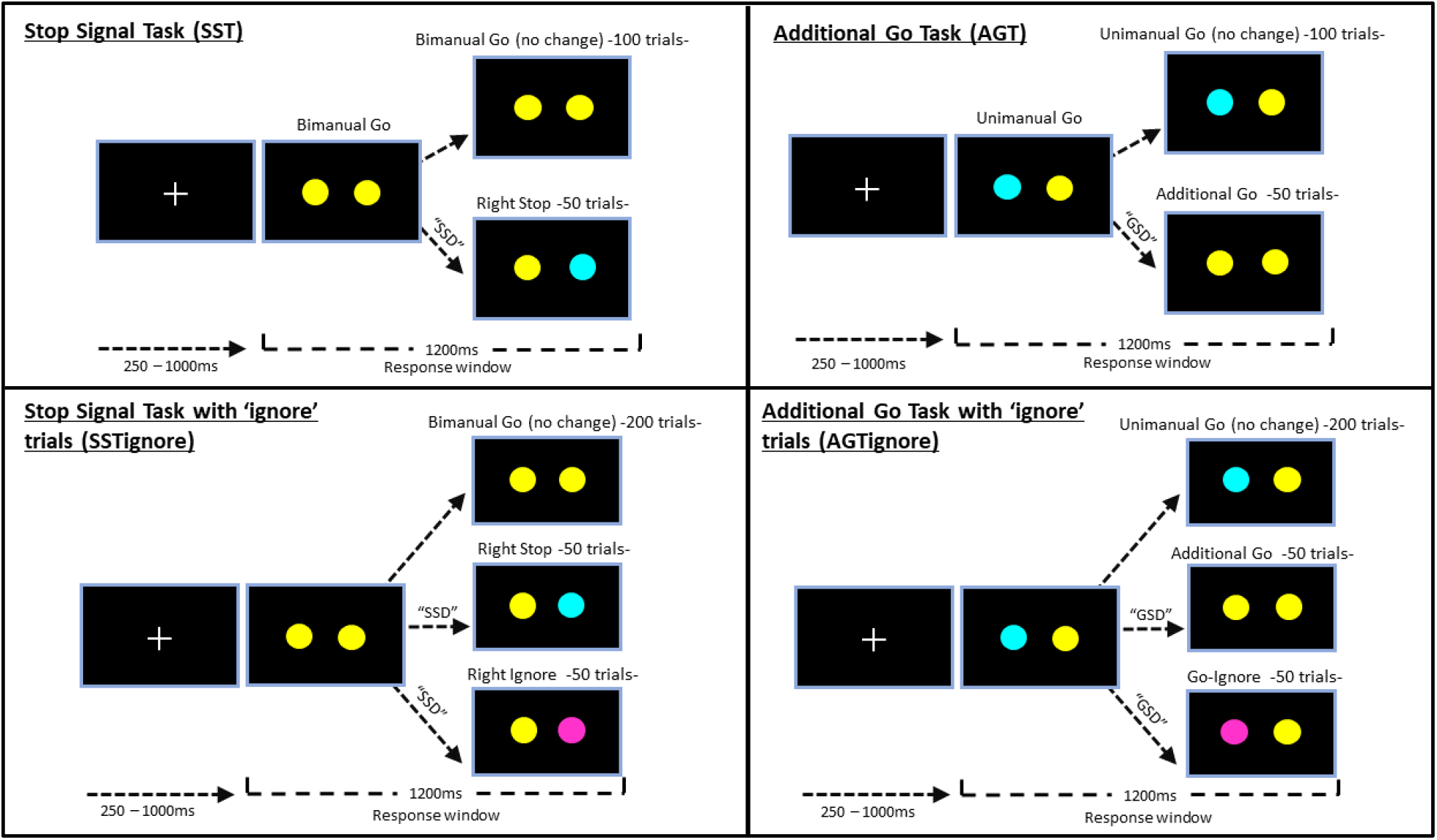
Trial types -and number of trials- in each of the four experimental conditions. The circles on the left and right were associated with responses with the left and right hands, respectively. Participants were required to respond to yellow circles, inhibit their responses when circles turned cyan, or maintain their initial response when circles turned magenta (i.e., ignore the colour change). Note SST and SSTignore left stop “catch trials” are not pictured. Unimanual go trials were evenly split between left go and right go trials. Additional go trials and go ignore trials were evenly split between trials in which the initial cue appeared on the left or right. Here, only trials in which the initial response was required on the right are pictured.

In each condition, trial order was pre-randomised, with the sole condition that no more than two of the infrequent trial types (all trial types other than “go” trials – see below) occurred consecutively. After every 50 trials, a short break was triggered. During the breaks, text appeared asking participants to rest their hands for a moment and to press one of the buttons when they were ready to continue. Below this text message, feedback was displayed indicating the proportion of correct responses and average reaction time (RT) of correct responses from the previous 50 trials.

#### 2.3.3. Stop Signal Task (SST)

The SST consisted of 153 trials (100 ‘bimanual go’ trials, 50 ‘right-stop’ trials and 3 ‘left-stop’ “catch” trials). During bimanual go trials, participants were required to press both the left and right button simultaneously when two yellow circles appeared (figure 1).

During right stop trials, after the presentation of the yellow circles, and following a variable ‘stop signal delay’ (SSD), the circle on the right would change colours from yellow to cyan. This colour change acted as the ‘stop signal’ and indicated that the participant should attempt to cancel their right-hand response (while continuing their left-hand response). A trial was considered correct when participants pressed *only* the left button (that is, they successfully inhibited only the right response) and the feedback screen would display “Good”. Any other response resulted in feedback displaying “incorrect response”. The three left-stop (catch) trials were included to ensure that the participant could stop the other hand and was responding to the location of the stop cue, rather than simply responding by stopping the right hand, irrespective of the cue’s location.

The SSD refers to the time between the presentation of the ‘go cue’ (the yellow circles) and the ‘stop cue’ (colour change to cyan). Initially, the SSD was set to 200ms, and this was adjusted using a staircase algorithm based on success in the previous trial. Following a correct right stop trial the SSD would increase by 50ms, making it harder to stop on the next stop trial. Following a failed right stop trial the SSD would decrease by 50ms, making it easier to stop on the next trial. Using this staircase procedure, the probability of successfully stopping approximates 50%, enabling assessment of both successful and failed stop trials, as well as calculation of SSRT (Verbruggen et al., 2019). The minimum and maximum SSDs were 50ms and 400ms, respectively.

During bimanual go trials, if participants reacted more than 150ms slower than their average RT in the baseline go block, text reading “You’ve slowed down” would be displayed during the feedback for that trial. This served to minimise the degree to which participants would slow their responses in anticipation of stop signals, a behavioural adjustment which can interfere with the staircase algorithm resulting in very high accuracy rates and invalidating further SSRT calculations (Verbruggen et al., 2019).

#### 2.3.4. SSTignore

The above SST task can be described as a ‘motor selective’ or ‘response selective’ SST (i.e., participants were required to inhibit only one component of a multi-component motor response – Wadsley et al., 2022). The ‘SSTignore’ task built upon this by combining motor selectivity with ‘*stimulus* selectivity’ (see section 1.3; Bissett & Logan, 2014). Accordingly, this task followed the same procedure as the SST, with the addition of ‘ignore’ trials, in which the right circle would turn magenta rather than cyan. When this occurred, participants were required to ignore the colour change and continue with the button press (i.e., respond bimanually as though the circle had remained yellow). This condition comprised of 200 bimanual go trials, 50 right stop trials (colour change to cyan), 50 right ignore trials (colour change to magenta) and 6 left stop catch trials (left circle turns cyan). Thus, the proportion of trials involving unexpected cues (stop or ignore cues) remained consistent with that in the SST (where 33% of trials were stop trials). The timing of the ignore cues was determined from the SSD value that was set following the previous stop trial. Performance on ignore trials did not influence SSD. As in the SST, if bimanual go trial reaction times were more than 150ms slower than that subject’s average RT in the baseline block, text reading “You’ve slowed down” would be displayed during the feedback for that trial.

#### 2.3.5. Additional Go Task (AGT)

The AGT was a choice reaction time task with an additional response required on 33% of trials. It featured 150 trials comprised of 100 unimanual go trials and 50 ‘additional go’ trials. Upon stimulus presentation in unimanual go trials one circle would be yellow and the other would be cyan. Participants were required to press *only* the button on the same side as the yellow circle (and refrain from pressing the button on the side of the cyan circle). Thus, in contrast to the SST and SSTignore, where no decision needed to be made regarding which side to respond on (because both effectors were involved in a go trial), the AGT essentially constituted a choice reaction time task. Go trials were split evenly between left-go and right-go trials (50% each). In the additional go trials the circle which was initially cyan would turn yellow after a variable delay; when this occurred, participants were required to press the corresponding button as quickly as possible. The delay (deemed ‘go signal delay’; GSD) was 200ms, 150ms, 100ms, 50ms, or 0ms (in which case both circles were yellow at onset of the stimuli). Of the 50 additional go trials there were 10 trials at each GSD, with 5 of these for each hand (i.e., 5 in which the additional cue occurred on the left following an initial right go stimulus, and 5 in which it occurred on the right following an initial left go stimulus). There was no requirement for the button presses on the additional go trials to be simultaneous; indeed, the initial (unilateral) choice response may (depending on the GSD) have been completed prior to the additional go stimulus occurring. However, we sought to determine if – when the GSD was presented close to action initiation of the first press – whether any RT delays on the initial response were observed due to presentation of an unexpected (additional go) stimulus.

Trial-wise feedback for correct responses in this task involved a single RT display for go trials and both RTs side by side (left response on the left, right response on the right) for additional go trials. Incorrect trials (where the wrong buttons were pressed, or no buttons were pressed) resulted in the presentation of “incorrect response” during the feedback screen. Unlike the SSTs, this task did not include “you’ve slowed down” warnings, as we did not anticipate that behavioural slowing would be an issue when participants were not anticipating stop signals.

#### 2.3.6. AGTignore

The AGTignore condition constituted 300 trials (200 unimanual go, 50 additional go, and 50 go-ignore trials). As such, the proportion of trials involving unexpected cues (additional go and go-ignore trials) remained consistent with all other tasks (33%). The unimanual go and additional go trials followed the same procedure as those trials in the AGT task described above. During go-ignore trials the cyan no-go stimulus changed colour to magenta. When this occurred, participants were required to behave as if there was no colour change (continue to refrain from responding on that side) and continue with the initial unilateral response. The delay of the ignore signal, like those of the additional go trials, was either 200ms, 150ms, 100ms, 50ms, or 0ms (started as magenta), with the ignore cues in half of the trials occurring on the left, and half on the right. Thus, in total each participant completed 5 ignore trials at each possible delay on both their left and right side.

### 2.4. Behavioural analysis

#### 2.4.1. Data processing

Prior to analysis, bimanual go trials in SST and SSTignore conditions were excluded if they were asynchronous (difference between left- and right-hand RTs of >50ms; 3.7% of bimanual go trials) or incorrect (i.e., the participant made a unimanual response, or made no response; 3.1% of all bimanual go trials). Furthermore, all trials with RTs <100ms, across all conditions, were excluded (<0.01%).

A generalised linear mixed model (GLMM) model approach was used for statistical analyses. All analyses were conducted using the statistical package Jamovi (The Jamovi Project, 2021) which runs using the statistical language R (R Core Team, 2021). GLMMs were run using the Jamovi module GAMLj (Gallucci, 2019). Analyses that were conducted on measures with reaction times (and the analysis conducted on ‘CancelTime’; see below) employed a gamma distribution with a log link function which accounts for positive skew often observed in RT data (Lo & Andrews, 2015). For each GLMM a maximal random effects structure was initially used. If this failed to converge a stepwise approach was taken to model simplification whereby interaction terms were first removed from the random effects structure, followed by slope terms (Barr et al., 2013). For the analyses on proactive slowing and interference effects (sections 2.4.2 and 2.4.3) participant ID was included as a random effects variable, allowing the use of trial-level RT data (rather than running an analysis on participant means). In contrast, the analysis run on SSRT uses a single score from each participant.

#### 2.4.2. Proactive slowing

The reactive component of action inhibition occurs following the environmental change which indicates that stopping is required (i.e., presentation of the stop signal). Prior to this, proactive inhibition may occur whereby adjustments are made within inhibitory networks to facilitate the potential upcoming action cancellation (Lavallee et al., 2014). A quantifiable behavioural component of proactive inhibition is proactive slowing (van de Laar et al., 2011). This is assessed by comparing participants’ RTs during an SST to their RTs outside of a stopping context (i.e., an equivalent response task but with no stop signals). Here, across each of the four conditions, RTs from correct responses in the initial Go-only block (see above) were compared to correct go trials from the main task of that condition (bimanual go trials in SSTs, unimanual go trials in AGTs). Our GLMM used the fixed factors of age group (younger, older), condition (SST, SSTignore, AGT, AGTIgnore), and block (go only block, main task). The final random effects structure included participant intercept and slopes of within subject variables (condition and block).

#### 2.4.3. Interference effects

The main analysis of interference effects included the presence/absence of EMG-recorded partial activations as a factor, so is presented below in section 2.5. For preliminary behavioural analyses of interference effects refer to supplementary materials.

It was theorised that the degree of interference observed in the AGT and AGTignore tasks may be contingent upon the precise timing of the additional cue, relative to the initial go cue. In order to test whether interference effects (from additional go cues) varied as a function of the timing of the additional go cue (GSD), analyses were run on data from the AGT and AGTignore conditions. Due to issues with model complexity and convergence, additional go trials were examined separately to go-ignore trials, and a separate analysis was run for each age group. In the analyses examining additional go trials a preliminary analysis revealed no effect of condition, so trials from from both the AGT and AGTignore were combined in one analysis. As such, four separate GLMMs were run: Additional go trials in the younger cohort, additional go trials in the older cohort, go-ignore trials in the younger cohort, and go-ignore trials in the older cohort. Unimanual go trials were included as a level of the GSD variable so that the six levels of the GSD variable were: No GSD (unimanual go), 0ms, 50ms, 100ms, 150ms, or 200ms. This then allowed us to test for significant interference effects at each of the possible GSDs via Bonferroni corrected post-hoc tests, using the unimanual go trials as the reference level.

#### 2.4.4. SSRT

SSRT was calculated using the integration method (Verbruggen et al., 2019). The horserace model which this method of SSRT calculation is based on requires that mean RTs on unsuccessful stops are faster than those on go trials. This requirement held true for all but one younger participant whose data (from both SST conditions) was removed from the SSRT analysis. Each participant has a single SSRT score for each condition. A general linear model was run on this data, with SSRT as the dependent variable and age and condition as independent variables.

### 2.5. EMG Analysis

#### 2.5.1. Data processing

EMG data processing was performed using the programming and numeric computing platform MATLAB (MathWorks, 2018). EMG signals were digitally filtered using a fourth-order band-pass Butterworth filter at 20–500 Hz. To detect the precise onset and offset times of EMG bursts we used a single-threshold algorithm (Hodges & Bui, 1996): Signals from each trial were rectified and smoothed by low-pass filtering at 50 Hz. Following this, a sliding window of 500 ms was used to find the segment with the lowest root mean squared (RMS) amplitude, which was used as baseline. EMG bursts were identified when the amplitude of the smoothed EMG was above 3 SD from baseline. For robustness, EMG bursts separated by less than 20 ms were merged together.

Given the detected onset and offset times, we used time constraints to identify two types of EMG burst: First, the *RT-generating burst* was identified as the last burst whose onset happened after the go signal and prior to the recorded button press. In addition, some trials exhibited *partial bursts*, i.e., EMG bursts that were cancelled before they were strong enough to generate an overt button press. These were identified in each hand as the earliest burst where (i) peak EMG occurred after the SSD/GSD and (ii) prior to the onset of the RT-generating burst from the responding hand; and (iii) peak EMG amplitude was greater than 10% of the average peak from that participant’s successful go trials (bimanual gos in SSTs and unimanual gos in AGTs). These time and amplitude constraints were critical to avoid inclusion of spurious bursts, in particular those initiated *after* the RT-generating burst, as these are likely related to mirror activity or other activity unrelated to the task. For each burst detected (RT-generating and partial), we extracted the times of onset, peak and offset.

To obtain EMG envelopes we used full-wave rectification and low-pass filtering at 10 Hz. EMG envelopes were used to extract the peak amplitude. For each subject, the amplitude of EMG envelopes from each hand was normalised by the average peak EMG from successful bimanual go trials in the SST and SSTignore, and unimanual go trials in the AGT and AGTignore.

#### 2.5.2. CancelTime (EMG-derived index of stopping latency)

On successful stop trials where a partial burst is observed, the latency of the peak amplitude of the partial EMG burst relative to that trial’s SSD can be used to determine a single trial estimate of stopping latency (Jana et al., 2020; Raud et al., 2022). That is, it is possible to calculate the delay between the appearance of the stop signal and the time at which muscle activity (that would have resulted in a button press had it not been cancelled) begins to decrease. In line with previous papers we refer to this as CancelTime (Jana et al., 2020).

Here, we have calculated CancelTime using trials in which partial activations were observed on the right hand, as this was the hand that was required to cancel its response in stop trials (while the left hand continued to respond). Prior to analysis, trials in which the peak of the partial burst occurred less than 100ms after the go signal were removed (1.3% removed). Outlier rejection had a lower cut-off of 50ms from SSD (Jana et al., 2020), removing < 2% of trials. No upper cut-off was included. A GLMM with a gamma distribution and a log link function was run on this trial-level data, using condition (SST, SSTignore) and age group (young, older) as factors. A maximal random effects structure was used.

#### 2.5.3. Interference effects

The interference effects observed in the preliminary behavioural analyses of AGT tasks were - albeit statistically significant - small compared to the interference effects in the SST and SSTignore tasks (see supplementary material). It was speculated that this difference in means does not wholly represent the actual degree of response slowing that occurs in these trials, but rather represents an aggregate of two types of responses: (i) Slower responses, in which the participant paused (and a more substantial interference effect would be observed), and (ii) faster responses, in which no interruption occurred. That is, if the additional stimulus was presented *after* a fast response had been executed, then there was no opportunity for pausing to occur in response to it. Thus, an analysis was run on the AGTignore task, including the presence/absence of a partial burst in the initially responding hand as a factor. After an initial model including age as a factor failed to converge, the analysis was re-run separately for older and younger cohorts. Both of these analyses used a random effects structure with participant intercepts and slopes for trial type and presence/absence of partial activation.

Similarly, an analysis including partial burst (present/absent) as a factor was also run on the SSTignore task. We focused on the left hand, as a response was always required on this hand (thus it provides a more consistent indicator of when motor output was interrupted). Both correct and incorrect stop and ignore trials were included in the analysis. This was done to compensate for the fact that trials with a partial burst were expected to be slower, and this facilitates a correct response on a stop trial, while an ignore trial will likely be classified as correct irrespective of whether a partial activation occurred. As with the analysis run on the AGTignore, an initial model, which also included age as a factor, failed to converge. Following this, the analysis was run separately for each age group. Both of these analyses used a random effects structure with participant intercepts and slopes for trial type and partial activation.

#### 2.5.4. Analyses of response amplitude

Based on stop-restart models of partial stopping (MacDonald et al., 2017), the amplitude of the continued response should be greater on trials where an already initiated go response is interrupted (as is observed in partial burst trials). To test for this, two additional GLMMs were run. On the data from the SST and SSTignore, right stop and right ignore trials were included in an analysis which used age, trial type (stop/ignore) and left-hand partial burst (present/not present) as independent variables. The dependent variable was peak amplitude of the left effector. The random effects structure included participant intercepts and slopes for both trial type and partial burst. On the data from the AGT and AGTignore additional go and go-ignore trials were included in an analysis which used age, trial type (additional go/go-ignore) and initial-hand partial burst (present/absent) as independent variables. After an initial random effects structure including all slopes demonstrated high singularity, a simplified random effects structure including only participant intercepts and slopes for partial burst was used.

## 3. Results

### 3.1. Stopping ability across the lifespan

#### 3.1.1. Accuracy

Trials success rates indicated that participants completed the task well in the go trials and ignore trials (success ∼93%-98%) and that staircasing of the stop signal realised stop success of ∼42-55%). A full table of trial success rates can be found in Appendix A. Descriptive results of behavioural RTs for each trial type for are also presented in Appendix A.

#### 3.1.2. Proactive slowing

The proactive slowing analysis revealed statistically significant main effects of condition *χ*^2^(3) = 10.93, *p* = 0.012, age group *χ*^2^(1) = 67.19, *p* < 0.001, and block *χ*^2^(1) = 53.22, *p* < 0.001. There was also a significant interaction effect between condition and block *χ*^2^(1) = 108.74, *p* < 0.001. The interactions between age and condition *χ*^2^(3) = 1.652, *p* = 0.647, and between age and block *χ*^2^(1) = 0.131, *p* = 0.717 were not statistically significant. The three way interaction was not statistically significant *χ*^2^(3) = 2.492, *p* = 0.477. In order to investigate the interaction effect between condition and block, a simple main effects analysis of block was conducted, split by condition (averaged across both age groups) This revealed statistically significant proactive slowing in the SST condition *χ*^2^(1) = 67.71, *p* < 0.001, and the SSTignore condition *χ*^2^(1) = 77.32, *p* < 0.001. No statistically significant proactive slowing was observed in the AGT condition *χ*^2^(1) = 0.195, *p* < 0.659. Surprisingly, the AGTignore condition revealed a statistically significant difference whereby responses were slightly *faster* in the main block compared to the go only block *χ*^2^(1) = 3.98, *p* < 0.046. Estimated marginal means are represented in table 1.

**Table 1.**
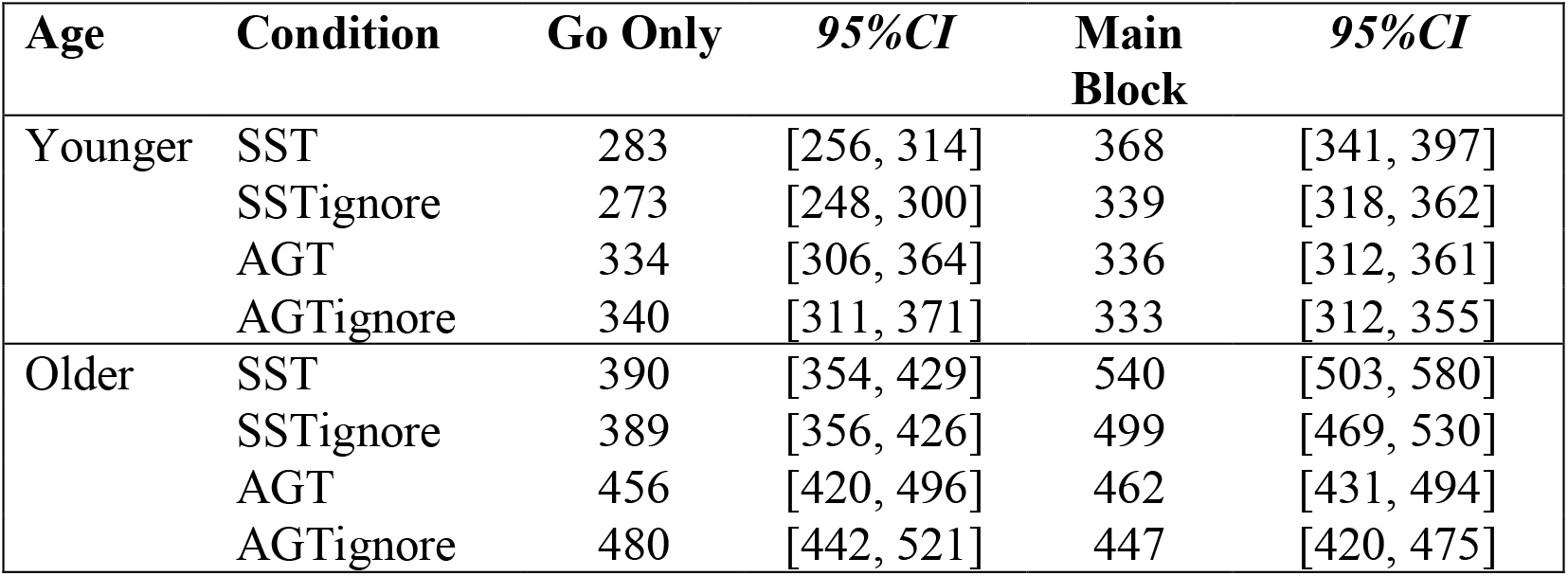
Estimated marginal means (in ms) from the proactive slowing analysis.

#### 3.1.3. SSRT and CancelTime

With respect to SSRT, the GLM revealed a statistically significant main effect of age, whereby older participants had slower SSRTs than younger adults *F*(1,88) = 94.96, *p* <0.001, *d* = 2.032. A statistically significant main effect of condition was also observed, whereby participants were slower at stopping in the SSTignore condition *F*(1,88) = 15.86, *p* <0.001, *d* = 0.830. The interaction between age and condition was not statistically significant *F*(1,88) = 1.71, *p* = 0.194. Mean SSRTs are reported in table 2.

**Table 2.** Mean SSRT and EMG measures of CancelTime for each age group in each stopping task (in ms)

| Condition | Age Group | SSRT | 95%CI | Canceltime | 95%CI |
| --- | --- | --- | --- | --- | --- |
| SST | Younger | 202 | [191, 214] | 164 | [149, 181] |
|  | Older | 270 | [258, 282] | 205 | [186, 227] |
| SSTIgnore | Younger | 234 | [223, 246] | 195 | [173, 220] |
|  | Older | 286 | [274, 298] | 241 | [213, 272] |

**Table 3:**
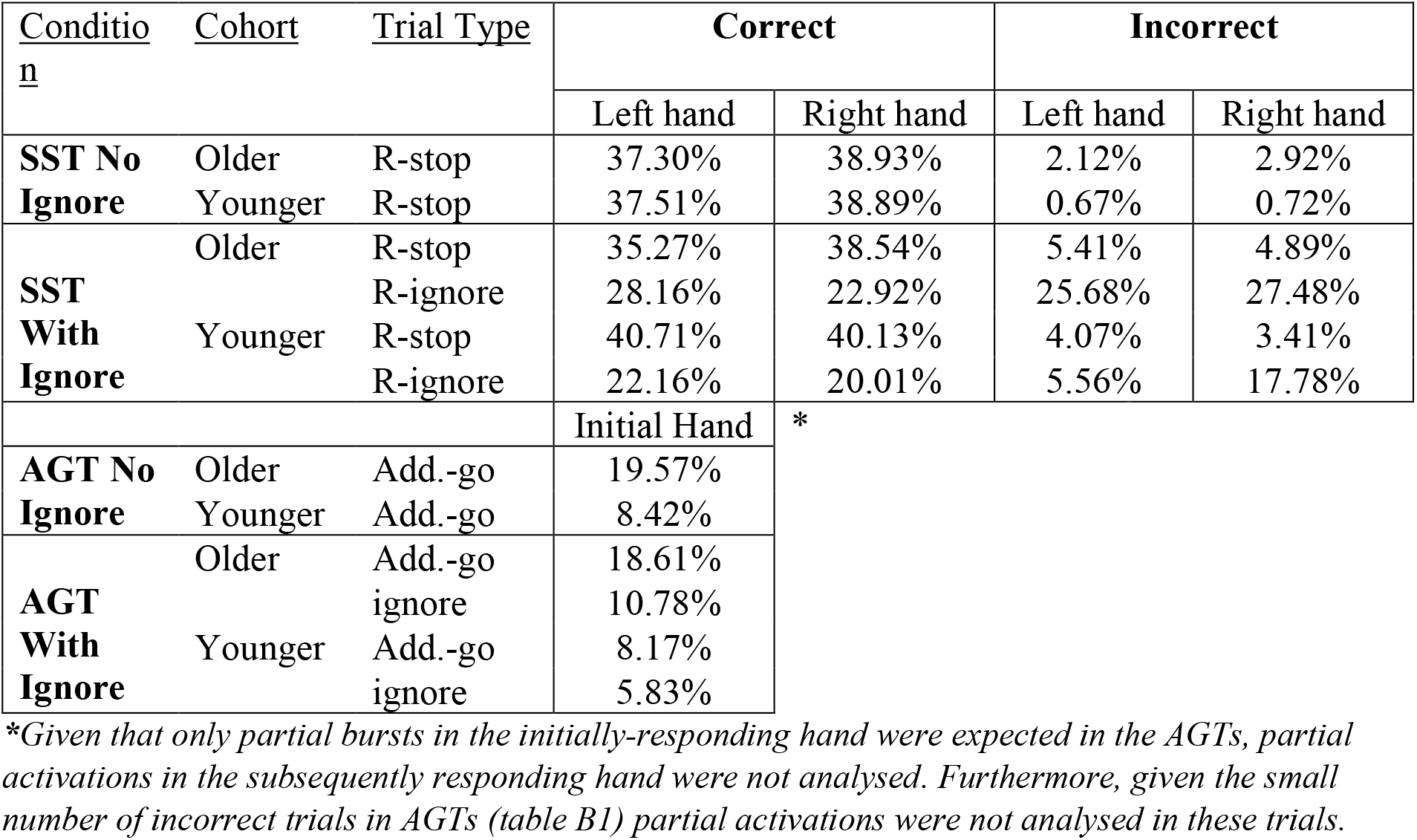
Percentage of Trials with Partial Activations.

With respect to CancelTime we observed a statistically significant main effect of age, whereby older participants had a longer CancelTime than younger participants, *χ*^2^(1) = 11.98, *p* <0.001, *d* = 0.45. Consistent with the results for SSRT, there was also a statistically significant main effect of condition, whereby CancelTime was longer in the SSTignore condition compared to the SST condition *χ*^2^(1) = 11.86, *p* < 0.001, *d* = 0.34. No statistically significant interaction between age and condition was observed *χ*^2^(1) = 0.03, *p* = 0.866. Mean CancelTimes are reported in table 2.

#### 3.1.4. Interference effects in AGTs split by go signal delay (GSD)

The analyses which examined interference effects in additional go trials and go-ignore trials at different GSDs are depicted in Figure 2a, and 2b, respectively, for both young and older adults.

**Figure 2:**
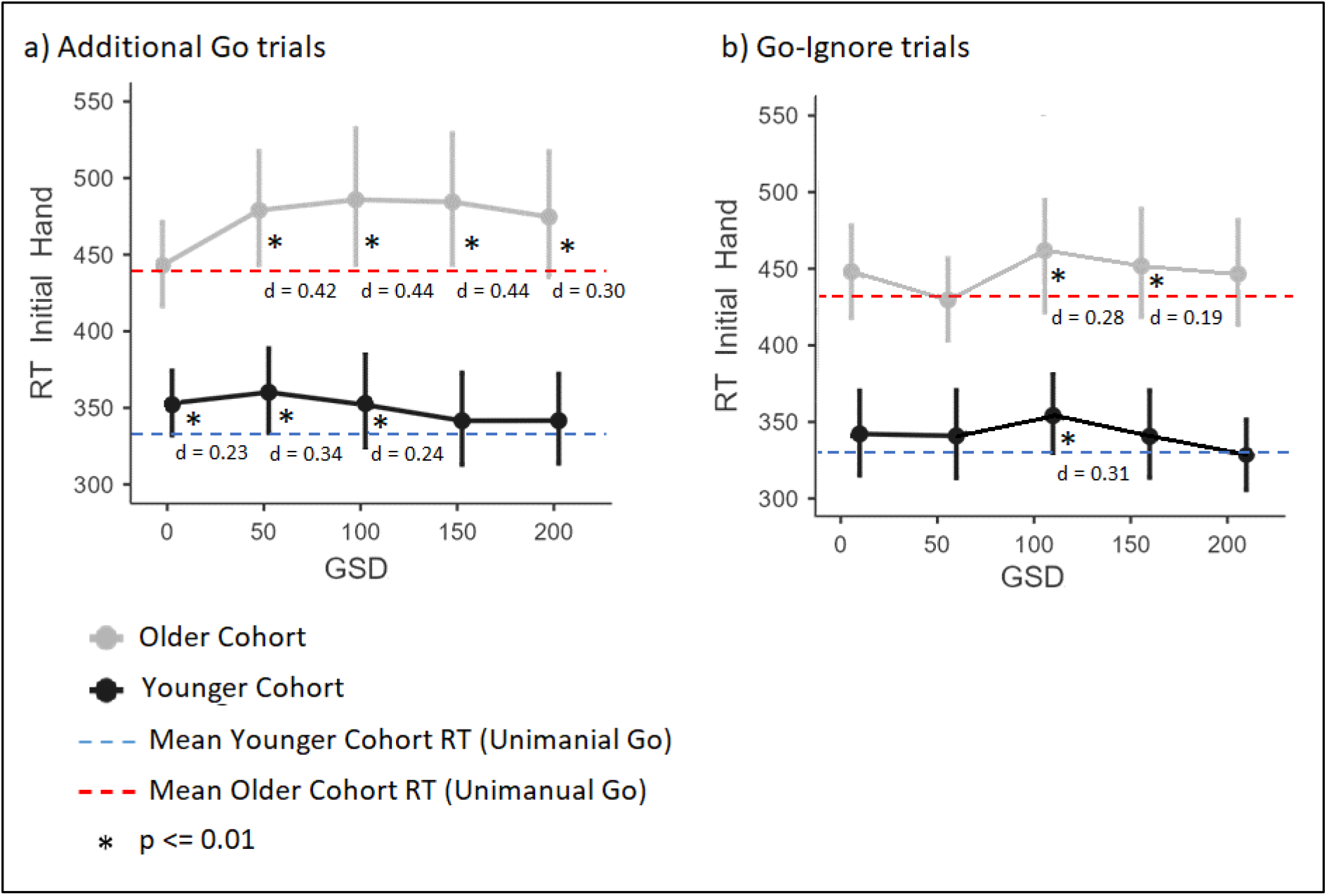
Estimated marginal means (RT in ms) of the initially responding hand for correct responses in additional go and go-ignore trials, split by go signal delay (GSD). Error bars represent 95%CIs from the four GLMMs that were run. Panel a) depicts additional go trials from both the AGT and AGTignore. Panel b) depicts go-ignore trials from the AGTignore. Mean RTs for unimanual responses are depicted for the younger (blue line) and older (red line) cohorts, and significant differences (at a Bonferroni corrected alpha level of p = 0.01) are indicated with *.

For the additional go trials (figure 2a), in the younger cohort a statistically significant main effect of GSD *χ*^2^(5) = 31.85, *p* <0.001 was observed. Post-hoc tests (using a Bonferroni corrected alpha level of p = 0.01) revealed the interference effect at GSDs of 0ms just failed to reach the adjusted a-priori alpha level (*z* = 2.56, *p* = 0.010). Statistically significant interference effects were observed at GSDs of 50ms (*z* = 4.24, *p* < 0.001), and 100ms (*z* = 2.91, *p* = 0.004). Interference effect at the GSDs of 150ms (*z* = 1.61, *p* = 0.107) or 200ms (*z* = 1.96, *p* = 0.050) did not reach the adjusted level of statistical significance. In the older cohort we observed a statistically significant main effect of GSD *χ*^2^(5) = 37.53, *p* <0.001. There was no statistically significant interference effect at a GSD of 0ms (*z* = 0.38, *p* = 0.707). However, statistically significant interference effects were observed at 50ms (*z* = 4.85, *p* < 0.001), 100ms (*z* = 4.63, *p* < 0.001), 150ms (*z* = 4.52, *p* < 0.001) and 200ms (*z* = 3.09, *p* = 0.002).

For go-ignore trials split by GSDs (Figure 2b), in the younger cohort we observed a statistically significant main effect of GSD *χ*^2^(5) = 16.96, *p* <0.001. Post hoc tests revealed interference effect at GSDs of 0ms (*z* = 1.87, *p* = 0.061) or 50ms (*z* = 2.03, *p* = 0.042) did not reach the adjusted level of significance. A significant interference effect was observed at 100ms (*z* = 3.44, *p* = 0.008). Interference effects at the GSDs of 150ms (*z* = 1.68, *p* = 0.093) or 200ms (*z* = 0.22, *p* = 0.823) were not statistically significant. For the older cohort we observed a statistically significant main effect of GSD *χ*^2^(5) = 26.14, *p* <0.001. The interference effect at a GSD of 0ms (*z* = 2.21, *p* = 0.027) and 50ms (*z* = 0.45, *p* < 0.653) were not statistically significant. There were statistically significant interference effects at 100ms (*z* = 2.92, *p* = 0.004) and 150ms (*z* = 2.72, *p* = 0.007), but not at 200ms (*z* = 1.53, *p* = 0.125).

### 3.2. Interference effects

#### 3.2.1 EMG profiles

Figures 3 and 4 represent the EMG profiles for the SSTignore task for the younger and older cohorts respectively while figures 5 and 6 represent the EMG profiles for the AGTignore task for the younger and older cohorts, respectively. All EMG profiles are synchronised to the onset of the RT generating burst. This allows for clear observation of the partial activations in stop and ignore trials (blue line). Moving the synchronisation reference away from the RT generating burst causes the shape and timing of the EMG profiles to “blur” due to the averaging of profiles whose peaks are not perfectly aligned in time. For this reason, the mean peak EMG in these figures appears to be slightly below 1.

**Figure 3:**
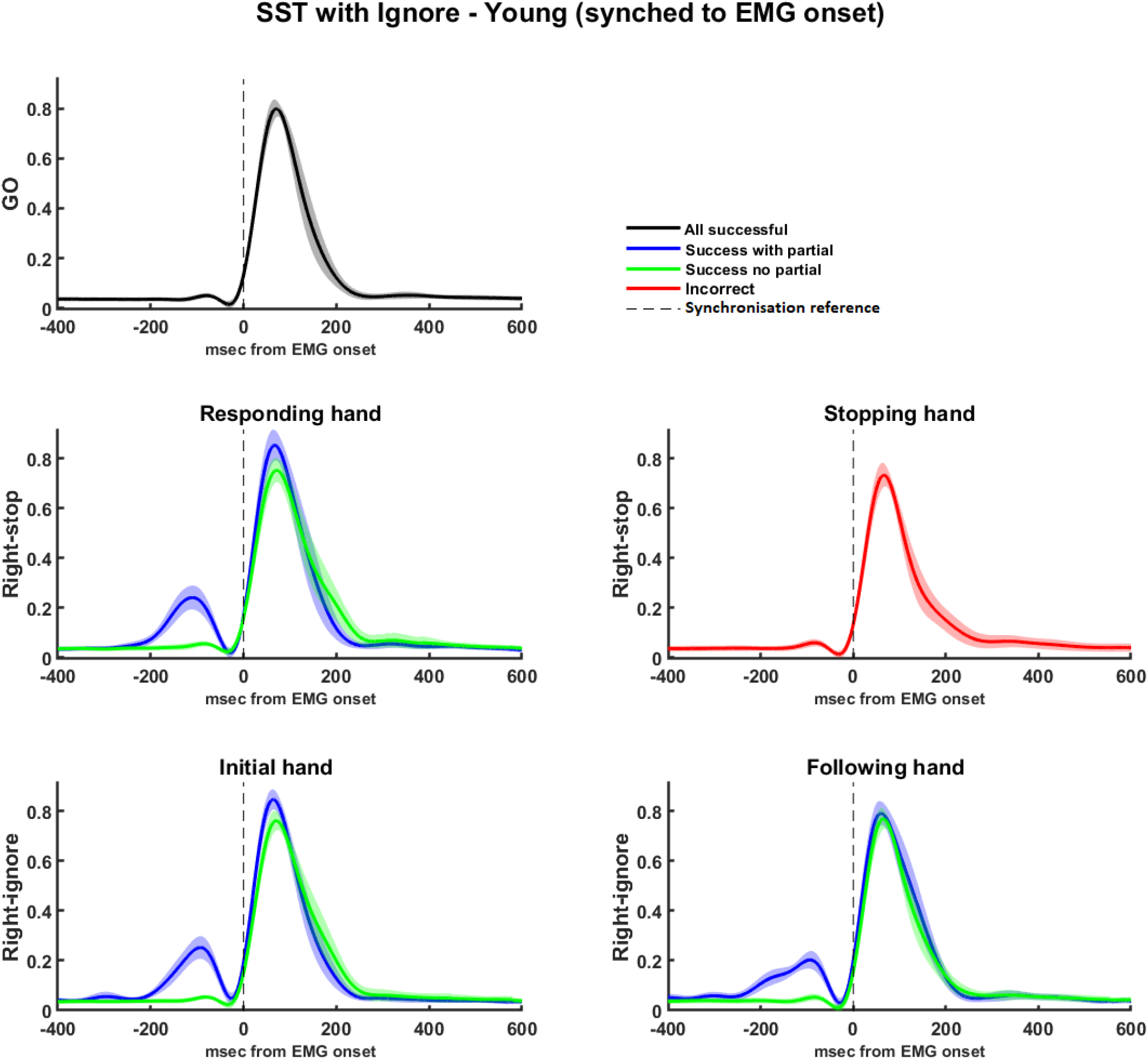
EMG profiles for the SSTignore for the younger cohort, split by hand, trial type, and presence/absence of partial burst. Trials are synchronised to the onset of the RT generating burst, in order to allow for observation of the presence of partial activations (and the subsequent inhibition). These are most notable on the left hand for right stop and right ignore trials and are represented with the blue line. Shaded areas represent 95%CIs.

**Figure 4:**
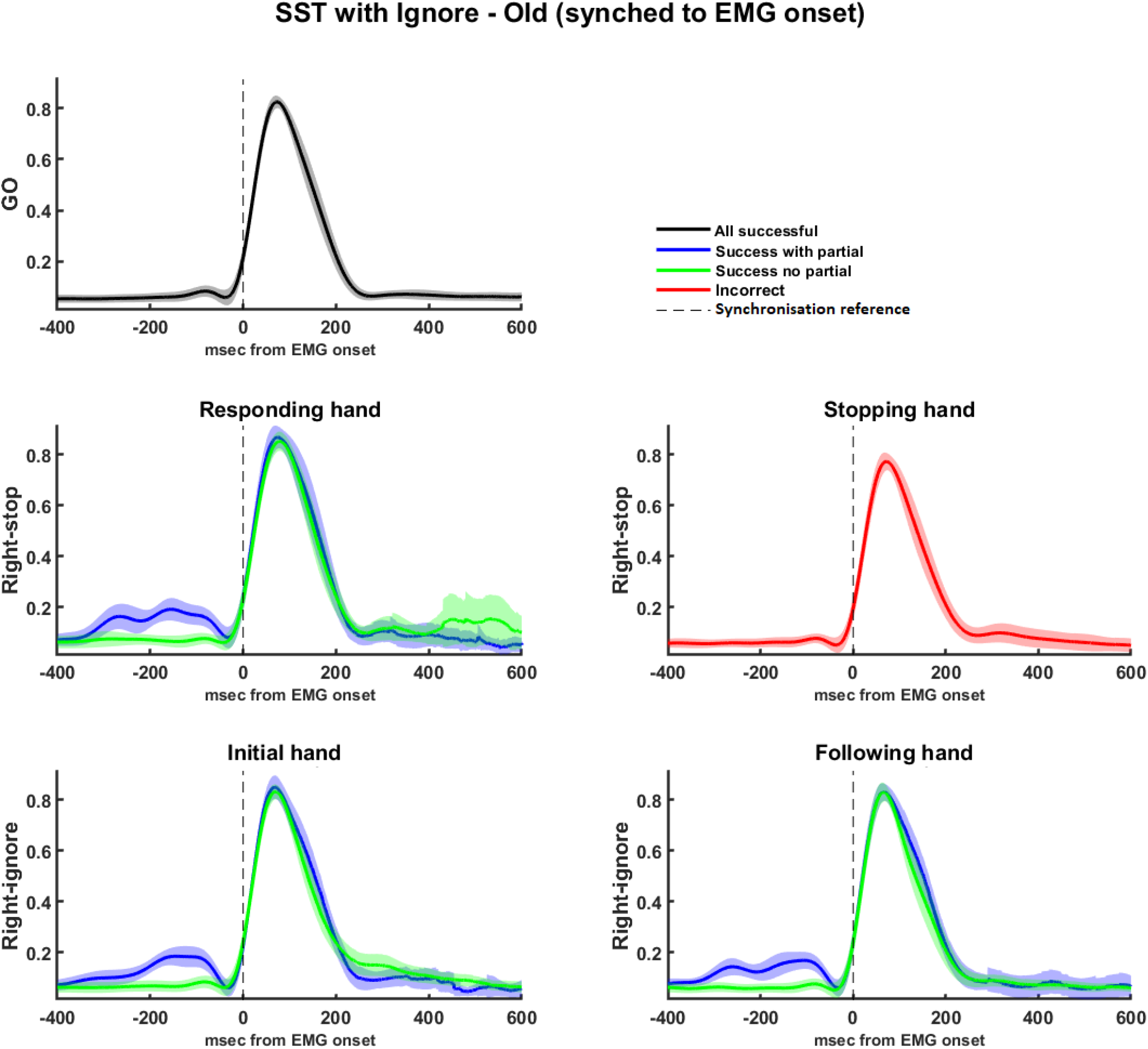
EMG profiles for the SSTignore for the older cohort, split by hand, trial type, and presence/absence of partial burst. Trials are synchronised to the onset of the RT generating burst, in order to allow for observation of the presence of partial activations (and the subsequent inhibition). These are most notable on the left hand for right stop and right ignore trials and are represented with the blue line. Shaded areas represent 95%CIs.

**Figure 5:**
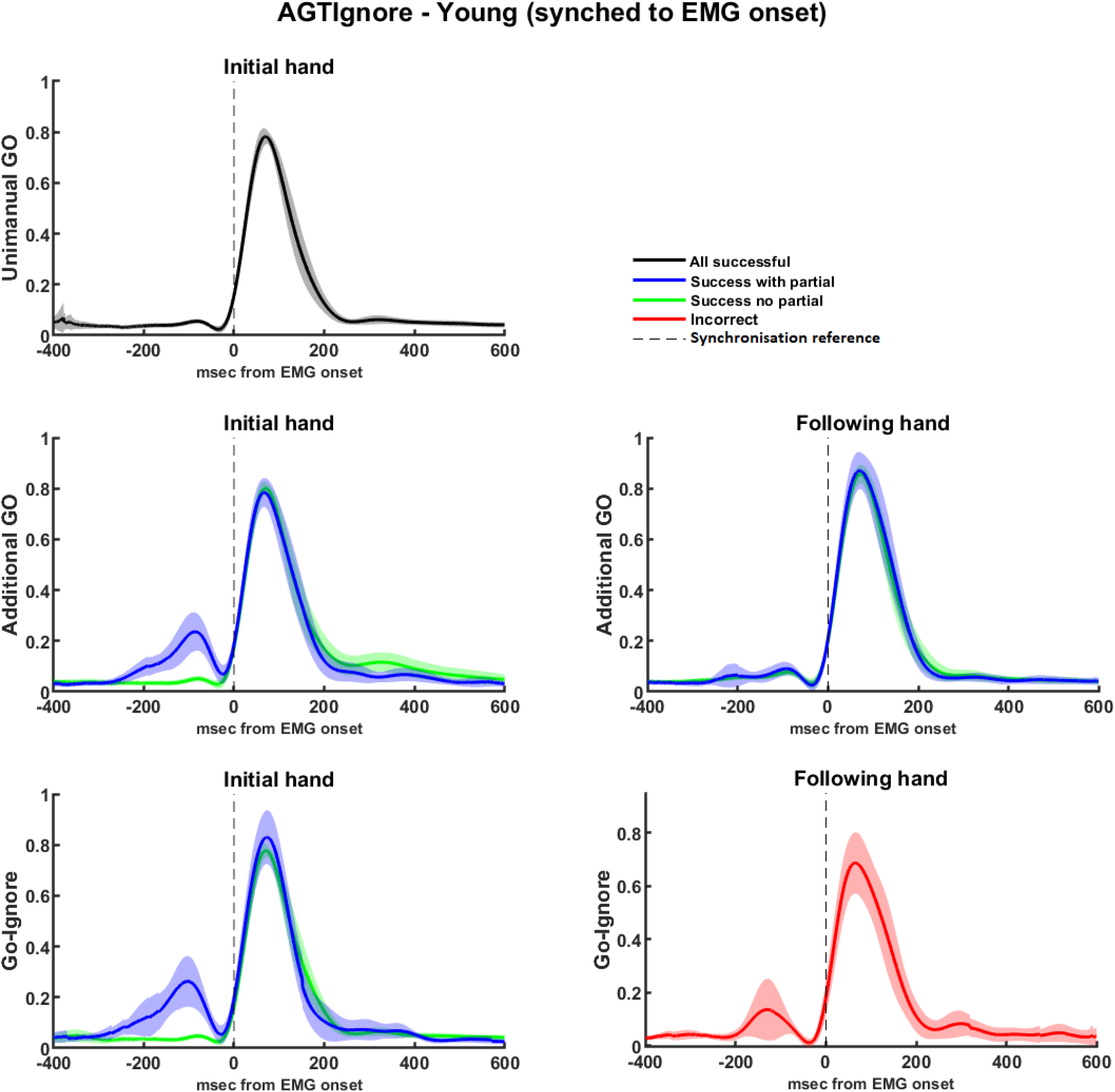
EMG profiles for the AGTignore for the younger cohort, split by hand, trial type, and presence/absence of partial burst. Trials are synchronised to the onset of the RT generating burst, in order to allow for observation of the presence of partial activations (and the subsequent inhibition). These are observable on the left side of left-then-bimanual trials, and the right side of right-then-bimanual trials. Shaded areas represent 95%CIs. Note that for successful go-ignore trials, the following hand has no RT-generating burst; hence, no EMG profile can be plotted when synchonised to the RT-generating burst. Here, incorrect trials are depicted synchronised to the RT generating burst in that hand.

**Figure 6:**
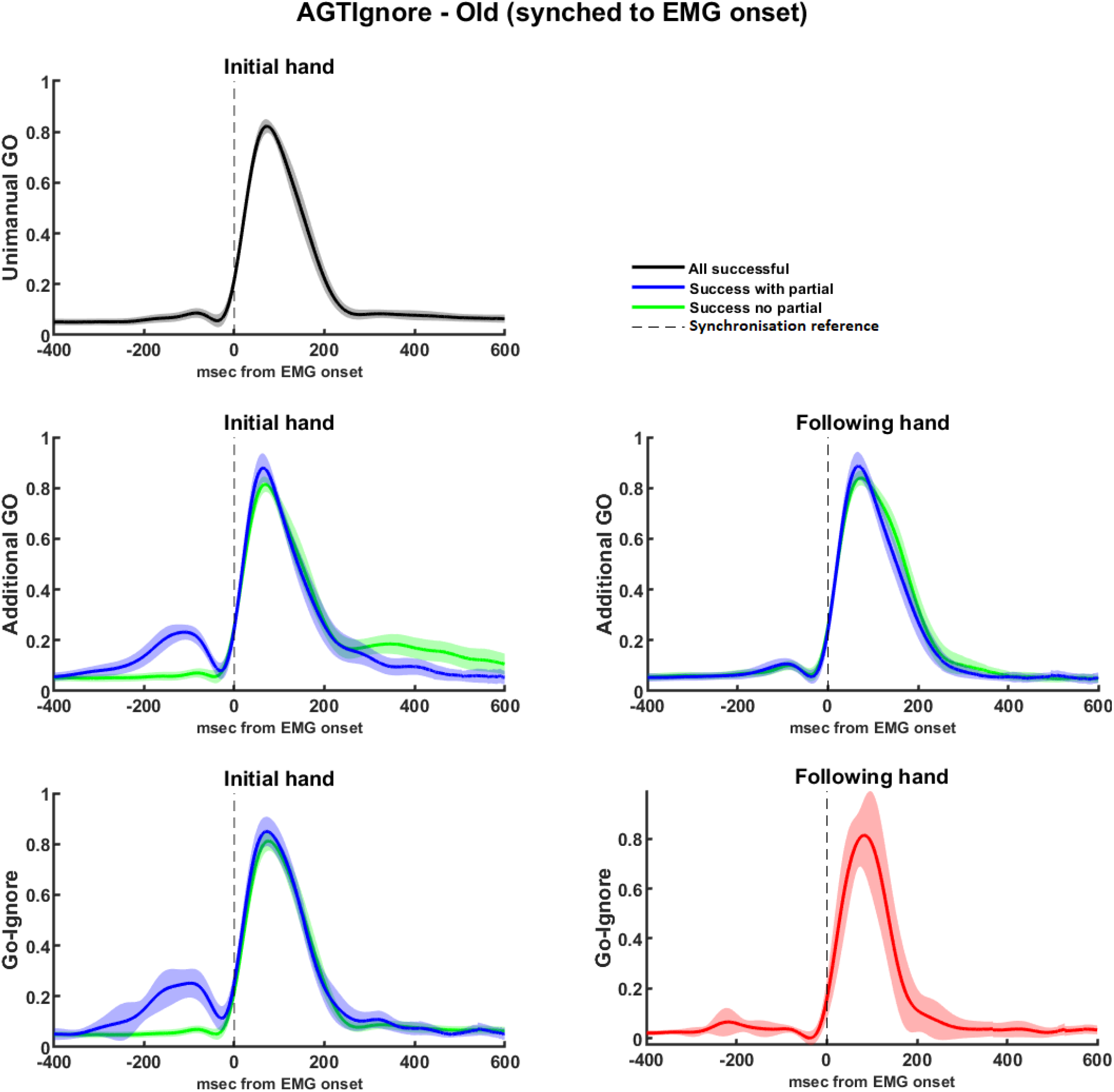
EMG profiles for the AGTignore for the older cohort, split by hand, trial type, and presence/absence of partial burst. Trials are synchronised to the onset of the RT generating burst, in order to allow for observation of the presence of partial activations (and the subsequent inhibition). These are observable on the left side of left-then-bimanual trials, and the right side of right-then-bimanual trials. Shaded areas represent 95%CIs. Note that for successful go-ignore trials, the following hand has no RT-generating burst; hence, no EMG profile can be plotted when synchonised to the RT-generating burst. Here, incorrect trials are depicted synchronised to the RT generating burst in that hand.

#### 3.2.2. Interference effects in SST and SSTignore with/without partial activations

Figure 7a depicts the results of the RT analysis run on the younger cohort. A significant main effect of trial type *χ*^2^(2) = 129.99, *p* <0.001, and partial burst was observed *χ*^2^(1) = 54.19, *p* <0.001. There was also a significant interaction effect between these two factors *χ*^2^(1) = 67.76, *p* <0.001. Bonferroni adjusted post-hoc tests revealed significant differences in RT between stop trials with (495ms) and without (443ms) partial activations (*z* = 3.24, *p* = 0.017, *d* = 0.50), as well as between ignore trials with (484ms) and without (361ms) partial activations (*z* = 8.97, *p* < 0.001, *d* = 1.10). Furthermore, stop and ignore trials both with and without partial activations were all significantly slower than go responses (340ms, *p* < 0.001). Notably, no significant difference was observed between stop and ignore trial RTs with partial activations (*z* = 0.94, *p* = 1.00).

**Figure 7:**
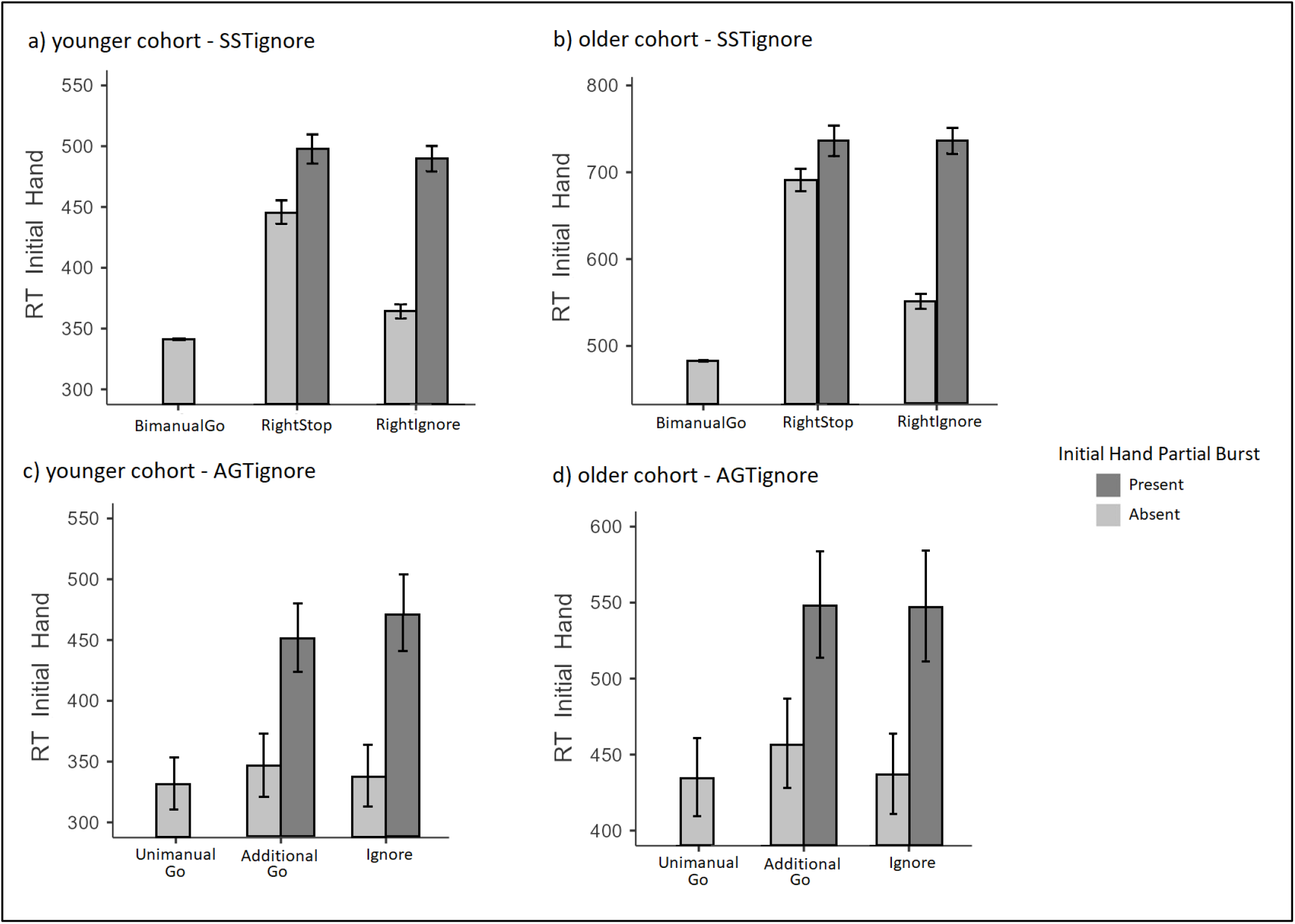
RT interference effects split by presence/absence of partial burst in the initially responding hand. The top two panels represent the SSTignore while the bottom two panels represent the AGTignore. The left two panels represent the younger cohort while the right two panels represent the older cohort. RTs presented in milliseconds. Error bars represent 95%CIs.

Figure 7b depicts the results of the analysis run on the older cohort. A significant main effect of trial type *χ*^2^(2) = 228.79, *p* <0.001, and partial burst was observed *χ*^2^(1) = 95.34, *p* <0.001. There was also a significant interaction effect between these two factors *χ*^2^(1) = 71.44, *p* <0.001. Bonferroni adjusted post-hoc tests revealed significant differences in RT between stop trials with (737ms) and without (665ms) partial activations (*z* = 3.71, *p* = 0.003, *d* = 0.47), as well as differences in RT between ignore trials with (735ms) and without (546ms) partial activations (*z* = 11.69, *p* < 0.001, *d* = 1.06). Furthermore, stop and ignore trials both with and without partial activations were all significantly slower than go responses (479ms, *p* < 0.001). As with the younger cohort, no significant difference was observed between stop and ignore trial RTs with partial activations (*z* = 0.116, *p* = 1.00).

#### 3.2.3 Interference effects in AGT and AGTignore with/without partial activations

Figure 7c depicts the results of the analysis run on the younger cohort. Overall, the main effect of trial type *χ*^2^(2) = 2.98, *p* = 0.225 was not statistically significant. However, there was both a significant main effect of partial burst *χ*^2^(1) = 67.92, *p* <0.001 and a significant interaction effect between these two factors *χ*^2^(1) = 5.214, *p* = 0.022. Bonferroni adjusted post-hoc tests revealed significant differences in RT between additional go trials with (451ms) and without (346ms) partial activations (*z* = 6.63, *p* < 0.001, *d* = 1.02), as well as differences in RT between ignore trials with (471ms) and without (337ms) partial activations (*z* = 8.27, *p* < 0.001, *d* = 1.30). Furthermore, RT of unimanual go trials (331ms) was significantly faster than all other trial types (*p* <0.001), except for ignore trials without partial activations observed (*z* = 1.73, *p* = 1.00). Notably, there was also no significant difference in RT between additional go and ignore trials with partial activations (*z* = 0.94, *p* = 1.00).

For the older cohort (Figure 7d) the main effect of trial type, *χ*^2^(2) = 3.22, *p* = 0.20 was not statistically significant, although there was a significant main effect of partial activations *χ*^2^(1) = 81.36, *p* < 0.001, whereby trials with partial activations had slower responses (560ms) than those without (440ms). Interestingly, the interaction effect between these two variables *χ*^2^(1) = 3.34, *p* = 0.068 did not reach statistical significance, suggesting that the effect partial activations had in slowing RT did not vary between conditions.

#### 3.2.4 Response amplitude split by presence/absence of partial burst

The results of the response amplitude analyses are depicted in Figure 8. The response amplitude analysis which included partial activations as a factor in the SST and SSTignore tasks revealed a statistically significant main effect of partial activation whereby trials with a prior partial burst (*M* = 106.89%, *SD* = 25.65%) exhibited a greater response amplitude burst (in the RT generating burst) than those without prior partial bursts (*M* = 101.29%, *SD* = 23.66%), *F*(1,47) = 30.65, *p* < 0.001. There was also a main effect of age, whereby older participants exhibited greater response amplitude (*M* = 105.72%, *SD* = 25.10%) than younger participants (*M* = 102.46%, *SD* = 23.75%), *F*(1,47) = 4.07, *p* = 0.049 and a statistically significant main effect of trial type, whereby stop trials (*M* = 105.75%, *SD* = 24.83%) resulted in greater response amplitude than ignore trials (*M* = 102.43%, *SD* = 23.94%), *F*(1,52) = 8.61, *p* = 0.005. The two way interactions between age and trial type *F*(1,52) = 0.00, *p* = 0.992, age and partial burst *F*(1,47) = 0.06, *p* = 0.805 and trial type and partial burst *F*(1,3878) = 0.62, *p* = 0.430 were all not statistically significant, as was the three-way interaction between age, trial type and partial burst *F*(1,3878) = 0.79, *p* = 0.376.

**Figure 8:**
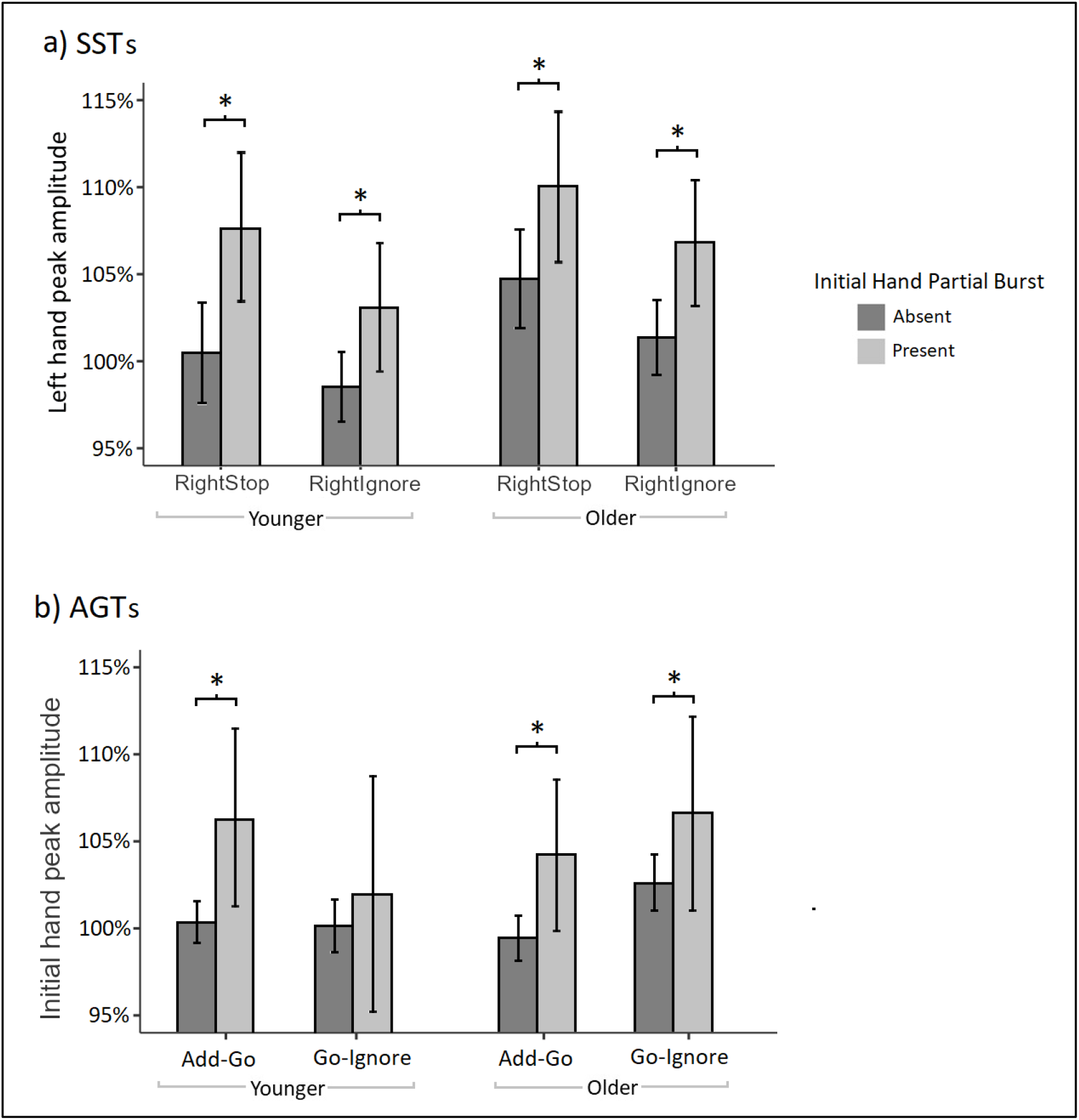
Panel a) represents the response amplitude of the non-cancelled (left) hand split by presence/absence of partial burst in the SST and the SSTignore tasks (pooled data). Panel b) represents the response amplitude of the initially cued hand split by presence/absence of partial burst in the AGT and AGTignore tasks (pooled data). Amplitudes are referenced as a percentage of that participant’s average peak EMG levels across all correct go trials in that condition (i.e., amplitude of the unimanual go trials in the AGTs and bimanual go trials in the SSTs are presented as 100% in panel a and b, respectively). Error bars represent 95%CIs. * = p < 0.05.

The response amplitude analysis which included partial activations as a factor in the AGT and AGTignore also revealed a statistically significant main effect of partial activation whereby trials with a prior partial activation exhibited greater response amplitude (*M* = 104.79%, *SD* = 28.36%) than those without (*M* = 100.66%, *SD* = 22.83%), *F*(1,54) = 5.74, *p* = 0.020. The main effects of age *F*(1,51) = 0.34, *p* = 0.561, and trial type *F*(1,6443) = 0.05, *p* = 0.825 were not statistically significant. There was a statistically significant interaction effect between age and trial type *F*(1,6443) = 5.56, *p* = 0.018. A test of simple main effects revealed that this interaction was driven by the older cohort demonstrating significantly greater response amplitude in go-ignore trials (*M* = 104.62%, *SD* = 26.35%) than additional go trials (*M* = 101.84%, *SD* = 25.17%), *t*(6730) = 2.15, *p* = 0.03, while for the younger cohort response amplitude in go-ignore trials (*M* = 101.71%, *SD* = 20.92%) was not statistically significant different to that in additional go trials (*M* = 103.37%, *SD* = 21.88%), *t*(6141) = 1.34, *p* = 0.19. The two interactions between age and partial burst *F*(1,54) = 0.02, *p* = 0.892, and between trial type and partial burst *F*(1,6440) = 1.33, *p* = 0.248, and the three-way interaction between age, trial type and partial burst *F*(1,6440) = 0.61, *p* = 0.434 were all not statistically significant.

## 4. Discussion

The goals of this experiment were threefold. Firstly, we examined interference effects (i.e., action execution delays) in a combined stimulus- and motor-selective stop signal tasks, using EMG analyses to elucidate the extent to which global inhibition is associated with attentional capture. Secondly, we used novel behavioural paradigms to identify similar interference effects (or an involuntary “pause” response, associated with attentional capture) *outside* of a stopping context. Finally, we aimed to extend our understanding of how the various complex processes associated with action cancellation (attentional capture, action inhibition) vary between healthy younger and older adults.

### 4.1. “Stopping interference” effects are observable outside a stopping context

Consistent with previous literature (Claffey et al., 2010; Drummond et al., 2018; Ko & Miller, 2011, 2013; Lavallee et al., 2014; Majid et al., 2012, 2013; Raud & Huster, 2017), we observed reaction time delays in motor selective stopping tasks where – following an initial multicomponent response (bilateral button press) – one response had to be inhibited. This has been termed the stopping interference (SI) effect (For review, see Wadsley et al., 2022). Moreover, similar RT delays (or costs) were exhibited when instead of requiring action inhibition, unexpected cues appeared that simply had to be *ignored*. To our knowledge, only one previous study has combined stimulus and motor selective stopping to investigate interference effects to ignore signals (Ko & Miller, 2013). In contrast to our current findings, that study did not observe statistically significant interference effects in response to ignore cues, concluding that the RT delays in selective stopping are primarily attributable to action cancellation, with minimal contribution from attentional capture processes. Given the growing body of research demonstrating that inhibition occurs broadly to unexpected stimuli (Wessel & Aron, 2017), we suggest that the larger interference effects in the current task (in both selective stop trials and ignore trials) are due to the differences in presentation frequency of stop and ignore cue. Specifically, 1/3 of trials in current study constituted trails in which infrequent (stop or ignore) cues were presented. In contrast, Ko and Miller (2013) had 12% of trials as go trials with 40% and 48% trials as stop and “controlled go” (ignore) trials, such that the absence of significant interference effects could be driven by the fact the ignore cues were expected (i.e., occurred more frequently than the ‘standard’ initial go response). This explanation would be consistent with the notion that the magnitude of the RT delay varies depending on the expectedness of the cue (Wadsley et al., 2022).

Using EMG allowed us to partition selective stop trials and ignore trials according to whether a covert “partial” response (i.e., not resulting in an overt button press) was, or was not, detected *prior* to the overt button press response. This allowed us to explicitly identify trials where inhibition was present, as in these trials a movement was initiated and subsequently suppressed. Notably, partial responses were often detected when ignore cues were presented, especially in the older participants (Figures 3 and 4), providing direct evidence that motor suppression occurred in these trials. When examining only those trials in which a partial activation was observed, there was no significant difference between response times on successful selective stop trials and those trials in which participants successfully ignored the ignore cues (Figure 7). This suggests that that an initial non-selective inhibition took place, and the degree of associated behavioural slowing was similar for both stop and ignore trials.

A key novel finding of the current experiment was that similar RT delays were also observed to infrequent cues in a task where action cancellation was not part of the response set. That is, when the initial task required a unilateral response, and participants had to infrequently execute an additional response with the contralateral limb, the initial response was often suppressed before being reinitiated. EMG profiles provide direct evidence that motor suppression occurred in response to the additional infrequent cues in these conditions (Figures 5 and 6). Unsurprisingly, those trials which demonstrated inhibition in the form of partial responses also resulted in greater behavioural interference effects (Figure 7). A recent experiment using a Go-NoGo task observed response slowing in response to unexpected cues, irrespective of whether they were associated with an imperative to go or not go (Sebastian et al., 2021). During that task predictive warning cues informed participants of whether an upcoming cue was likely to be a go cue (requiring a button press) or a no-go cue (where no button press was required). On a subset of trials, this warning cue was misinformative (that is, it made an incorrect prediction). These trials were associated with activation of the hyperdirect pathway, observed by fMRI, along with behavioural slowing, irrespective of whether the cue that appeared was a go cue or a no-go cue. It seems feasible that the same inhibitory process was involved in the neurophysiological effects that are observed in response to infrequent additional go responses in the current task. Research combining a behavioural paradigm similar to that used in the current experiment with brain imaging techniques would help shed further light on this issue.

Our results also demonstrate interference effects with regards to response amplitude, with greater peak amplitudes in the continuing effector during selective stops relative to the same hand during bimanual go trials (Figure 8). This finding can be explained in the context of the activation threshold model of selective stopping (MacDonald et al., 2014; 2017) in which an initial nonselective inhibition occurs following the stop signal, raising the activation threshold that is required for a subsequent (unilateral) response to occur. As a result, when the action occurs, it occurs with greater force. Interestingly, across all trial types with additional cues (stop trials, ignore trials, additional go trials and go-ignore trials), responses which demonstrated inhibition in the form of a partial burst, resulted in greater response amplitude in the continued response (Figure 8)^1^. This suggests that the increase in amplitude that occurs following selective stops generalises to instances of inhibition outside of stopping contexts (MacDonald et al., 2017).

### 4.2. Inhibition outside of stopping contexts may represent a “pause” response

A recently proposed two-stage model of action inhibition posits that two complimentary inhibitory processes facilitate action stopping (Diesburg & Wessel, 2021; Schmidt & Berke, 2017). In their review, Diesburg and Wessel review several lines of research including human and animal studies and conclude that the evidence is consistent with a two-stage model of action stopping. It is further suggested that that these two processes may demarcate the difference between the inhibition associated with attentional capture (the *pause* process), and that which reflects the voluntary cancellation of an action (the *cancel* process). It is also suggested that the pause process manifests as global/non-selective suppression of motor output (i.e., also occurs in effectors not associated with the current task), while the cancel process can be consciously targeted to specific effectors. Notably, Diesburg and Wessel (2021) also propose that the pause process may be observable via EMG. In developing our novel task which involved infrequent cues requiring additional actions, rather than action cancelling, it was theorised that any observed inhibition (detected via delayed response times) could be unambiguously attributed to involuntary attentional orienting processes because the task demands *did not* require any action cancellation. Furthermore, participants were encouraged (via task instructions) to respond quickly, so any RT delays that occurred sat in opposition to task goals. Thus, it is plausible that the RT delays that were observed in the AGTs in the current experiment provide observable evidence of the pause process at the level of the muscle.

An alternative explanation of the interference effects observed in the AGTs is that they represent a decision-making process in relation to the functional coupling of the two effectors (Wadsley et al., 2019). Specifically, if the second cue occurs before a certain time point, there may be a natural tendency to delay the initial movement and combine the two distinct unimanual movements into a single bimanual motor plan. However, it should be noted that such a decision-making process would still require inhibition, to delay the unimanual movement such that a coupled bimanual action could occur.

### 4.3. Interference effects occur over a longer temporal window for older adults

In the SST and SSTignore, a staircase algorithm allows the presentation of the stop/ignore signal to occur at the cusp of where action cancellation is possible for a given participant. However, in the AGTs such staircasing was not possible because there was no success/failure parameter upon which to vary the presentation of the additional go stimuli (i.e., no matter how late an additional go stimuli appears, participants can always respond to it, albeit with greater delay relative to the initial unimanual go response). Accordingly, we used 5 preselected go signal delays (GSDs). Including GSD as a factor in the analysis revealed that the timing of the additional go cue influences the extent of the observed interference (Figure 2). Interestingly, interference effects were not observed in the younger cohort at GSDs of 150ms and 200ms, despite average behavioural responses (i.e., button presses) occurring at approximately 350ms. This suggests that once a certain point is passed in the movement planning and execution process, attentional capture (i.e., presentation of new infrequent stimuli) will not interfere with current motor plans. In contrast, interference effects were observed in older adults at GSDs of 150 and 200ms. This reflects the slower reaction times of the older cohort; however, it also suggests that the window in which attentional capture (and the inhibition associated with this) can interfere with current motor plans is longer in older adults.

### 4.4. Inhibitory performance across the lifespan

Reactive inhibition was indexed via the behavioural estimate of stopping performance (SSRT) and EMG observations of ‘CancelTime’ (Jana et al., 2020). Consistent with prior research, both these measures suggest slower inhibitory processes in older compared younger adults, in stopping tasks with, and without, ignore stimuli (Bloemendaal et al., 2016; Coxon et al., 2016; Hsieh & Lin, 2016; Kleerekooper et al., 2016; Smittenaar et al., 2015).

Both age groups demonstrated slower reactive inhibition during the SSTignore task compared to the regular SST task. This difference was observed with both SSRT and CancelTime. It is important to note that the frequency at which stop signals were presented was lower in the SSTignore task compared to the SST task (15% v. 30% of trials, respectively) which may have contributed to the observed increase in stopping latency in the SSTignore task (c.f. Bissett & Logan, 2014), although this finding is not universal – see (Ramautar et al., 2004). An alternate interpretation of this is that the additional processing involved in discriminating between stop and ignore stimuli requires time, increasing stopping latency. Interestingly, the effect of condition (SST vs SSTignore) did not significantly interact with age. That is, both older and younger adults demonstrated slower reactive inhibition in the more complex task, and the degree of slowing was comparable across the two age groups. Past research observes mixed results on this. For instance, a study by Hsieh & Lin (2017) reported that older adults demonstrated greater stopping deficits compared to younger adults when an SST required stimulus discrimination. Notably, that study used auditory stimulus discrimination, whereby participants were required to discriminate between a true stop signal (tone at 1000Hz) and a ‘false’ stop signal (500Hz). Given that discrimination of auditory stimuli degrades with age (Clinard et al., 2010), this may have been particularly challenging for older adults, exaggerating the apparent changes to reactive inhibition. Other research, using visual stop signals, failed to observe age-related differences in stopping performance when stimulus identification was required (van de Laar et al., 2011). Assessing stimulus selective SST performance using various sensory modalities in the one experiment may clarify whether age-declines are indeed driven by task complexity or sensory (auditory) discrimination per se. Nonetheless, in the current study, there was no evidence that the requirement of stimulus discrimination interfered with stopping ability for older adults to a greater degree than younger adults.

### 4.5. Conclusions

In conclusion, we have provided novel evidence of behavioural and physiological manifestations associated with reactive inhibition in two novel tasks in which behavioural imperatives to cancel movement are not present, extending our understanding of these mechanisms and elucidating their broad relevance across greater range of motor control/decision tasks. Our results provide further evidence that current indices of reactive inhibition in stop signal tasks capture a combined measure of both an involuntary suppression of motor activity associated with attentional capture (which is not solely associated with subsequent possibility of action cancellation) and the voluntary cancellation of an action.

## Declarations of interest

none

## Author contributions

**Simon Weber:** conceptualisation, methodology, software, writing – original draft, formal analysis, investigation. **Sauro Salomoni:** conceptualisation, methodology, software, formal analysis, writing – review & editing. **Callum Kilpatrick:** conceptualization, investigation. **Mark Hinder:** conceptualization, resources, writing – review & editing, supervision, project administration, funding acquisition.

## Appendix A

**Table A.1:** Percentage of successful trials

| **Condition** | **Cohort** | **Trial Type** | **% Correct** |
| --- | --- | --- | --- |
| **SST** | Older | Bimanual Go | 93.75% |
|  |  | Right Stop | 55.39% |
|  | Younger | Bimanual Go | 98.25% |
|  |  | Right Stop | 49.55% |
| **SSTignore** | Older | Bimanual Go | 95.69% |
|  |  | Right Stop | 51.30% |
|  |  | Right Ignore | 90.43% |
|  | Younger | Bimanual Go | 98.43% |
|  |  | Right Stop | 42.33% |
|  |  | Right Ignore | 97.33% |
| **AGT** | Older | Unimanual Go | 96.52% |
|  |  | Additional Go | 95.48% |
|  | Younger | Unimanual Go | 97.50% |
|  |  | Additional Go | 98.58% |
| **AGTignore** | Older | Unimanual Go | 97.30% |
|  |  | Additional Go | 93.57% |
|  |  | Go-Ignore | 97.04% |
|  | Younger | Unimanual Go | 98.19% |
|  |  | Additional Go | 97.91% |
|  |  | Go-Ignore | 95.33% |

**Table A.2:** Mean RTs (ms) of correct responses

| Condition | AgeGroup | TrialType | RT | 95%CI | Interference | *d* |
| --- | --- | --- | --- | --- | --- | --- |
| **SST** | Younger | Bimanual Go | 370 | [367 – 374] | - |  |
|  |  | Stop Right | 481 | [473 – 489] | 111 | 1.15 |
|  | Older | Bimanual Go | 524 | [518 – 530] | - |  |
|  |  | Stop Right | 687 | [676 – 698] | 163 | 1.19 |
| **SST Ignore** | Younger | Bimanual Go | 343 | [341 – 345] | - |  |
|  |  | Stop Right | 477 | [468 – 487] | 134 | 1.41 |
|  |  | Ignore Right | 392 | [385 – 399] | 49 | 0.48 |
|  | Older | Bimanual Go | 482 | [479 – 486] | - |  |
|  |  | Stop Right | 707 | [694 – 719] | 225 | 1.70 |
|  |  | Ignore Right | 604 | [592 – 617] | 122 | 0.75 |
| **AGT** | Younger | Unimanual Go | 341 | [337 – 344] | - |  |
|  |  | Additional Go | 355 | [349 – 360] | 14 | 0.15 |
|  | Older | Unimanual Go | 453 | [449 – 457] | - |  |
|  |  | Additional Go | 479 | [472 – 486] | 26 | 0.24 |
| **AGT Ignore** | Younger | Unimanual Go | 336 | [334 - 338] | - |  |
|  |  | Additional Go | 356 | [350 - 361] | 20 | 0.22 |
|  |  | Ignore Cue | 351 | [345 – 356] | 15 | 0.17 |
|  | Older | Unimanual Go | 437 | [435 – 440] | - |  |
|  |  | Additional Go | 468 | [461 – 475] | 31 | 0.34 |
|  |  | Ignore Cue | 452 | [446 – 459] | 15 | 0.16 |

*RTs in SST and SSTignore represent left hand reaction times. RTs in AGT and AGTignore conditions represent reaction times of the initially responding hand. 'Interference' represents the difference between the mean RTs observed in that trial type and those observed in that condition's 'go' response. For preliminary analyses of interference effects in behavioural data (without presence/absence of partial burst included) refer to supplementary material.*

## Footnotes

1 With the sole exception of younger participants in the go-ignore trials.

## Notes

Funding: This work was supported by the Australian Research Council Discovery program [DP200101696] and a University of Tasmania Graduate Research Scholarship.

### Competing Interest Statement

The authors have declared no competing interest.

## References

Aron, A. R. (2011). From reactive to proactive and selective control: developing a richer model for stopping inappropriate responses. Biological Psychiatry, 69(12), e55–e68. https://doi.org/10.1016/j.biopsych.2010.07.024.From

Aron, A. R., & Verbruggen, F. (2008). Stop the presses: Dissociating a selective from a global mechanism for stopping: Research article. Psychological Science, 19(11). https://doi.org/10.1111/j.1467-9280.2008.02216.x

Barr, D. J., Levy, R., Scheepers, C., & Tily, H. J. (2013). Random effects structure for confirmatory hypothesis testing: Keep it maximal. Journal of Memory and Language, 68(3), 255–278. https://doi.org/10.1016/J.JML.2012.11.001

Bissett, P. G., Jones, H. M., Poldrack, R. A., & Logan, G. D. (2021). Severe violations of independence in response inhibition tasks. Science Advances, 7(12), 4355–4372. https://doi.org/10.1126/SCIADV.ABF4355/SUPPL_FILE/ABF4355_SM.PDF

Bissett, P. G., & Logan, G. D. (2014). Selective stopping? Maybe not. Journal of Experimental Psychology: General, 143(1), 455–472. https://doi.org/10.1037/a0032122

Bloemendaal, M., Zandbelt, B., Wegman, J., van de Rest, O., Cools, R., & Aarts, E. (2016). Contrasting neural effects of aging on proactive and reactive response inhibition. Neurobiology of Aging, 46, 96–106. https://doi.org/10.1016/j.neurobiolaging.2016.06.007

Buchman, A. S., Wilson, R. S., Bienias, J. L., & Bennett, D. A. (2009). Change in Frailty and Risk of Death in Older Persons. Experimental Aging Research, 35(1), 61–82. https://doi.org/10.1080/03610730802545051

Cai, W., Oldenkamp, C. L., & Aron, A. R. (2011). A proactive mechanism for selective suppression of response tendencies. Journal of Neuroscience, 31(16), 5965–5969. https://doi.org/10.1523/JNEUROSCI.6292-10.2011

Cambridge Electronic Design Ltd. Signal 4 for Windows. https://ced.co.uk/products/signal

Chen, W., de Hemptinne, C., Miller, A. M., Leibbrand, M., Little, S. J., Lim, D. A., Larson, P. S., & Starr, P. A. (2020). Prefrontal-Subthalamic Hyperdirect Pathway Modulates Movement Inhibition in Humans. Neuron, 106(4). https://doi.org/10.1016/j.neuron.2020.02.012

Claffey, M. P., Sheldon, S., Stinear, C. M., Verbruggen, F., & Aron, A. R. (2010). Having a goal to stop action is associated with advance control of specific motor representations. Neuropsychologia, 48(2), 541–548. https://doi.org/10.1016/j.neuropsychologia.2009.10.015

Clinard, C. G., Tremblay, K. L., & Krishnan, A. R. (2010). Aging alters the perception and physiological representation of frequency: Evidence from human frequency-following response recordings. Hearing Research, 264(1–2), 48–55. https://doi.org/10.1016/J.HEARES.2009.11.010

Coxon, J. P., Goble, D. J., Leunissen, I., Van Impe, A., Wenderoth, N., & Swinnen, S. P. (2016). Functional Brain Activation Associated with Inhibitory Control Deficits in Older Adults. Cerebral Cortex, 26(1), 12–22. https://doi.org/10.1093/cercor/bhu165

Coxon, J. P., Stinear, C. M., & Byblow, W. D. (2007). Selective inhibition of movement. Journal of Neurophysiology, 97(3), 2480–2489. https://doi.org/10.1152/jn.01284.2006

Diesburg, D. A., & Wessel, J. R. (2021). The Pause-then-Cancel model of human action-stopping: Theoretical considerations and empirical evidence. Neuroscience and Biobehavioral Reviews, 129(July), 17–34. https://doi.org/10.1016/j.neubiorev.2021.07.019

Drummond, N. M., Cressman, E. K., & Carlsen, A. N. (2018). Increased response preparation overshadows neurophysiological evidence of proactive selective inhibition. Psychology and Neuroscience, 11(1), 1–17. https://doi.org/10.1037/pne0000130

Gallucci, M. (2019). GAMLj: General analyses for linear models. [jamovi module]. https://gamlj.github.io/.

González-Villar, A., Galdo-Álvarez, S., & Carrillo-de-la-Peña, M. T. (2022). Neural correlates of unpredictable Stop and non-Stop cues in overt and imagined execution. Psychophysiology, 59(7), 1–14. https://doi.org/10.1111/psyp.14019

Gulberti, A., Arndt, P. A., & Colonius, H. (2014). Stopping eyes and hands: Evidence for non-independence of stop and go processes and for a separation of central and peripheral inhibition. Frontiers in Human Neuroscience, 8(1 FEB), 61. https://doi.org/10.3389/FNHUM.2014.00061/BIBTEX

Hodges, B., & Bui, B. (1996). A comparison of computer-based methods for the determination of onset of muscle contraction using electromyography. Electroencephalography and Clinical Neurophysiology, 101(6), 511–519. https://doi.org/10.1016/S0013-4694(96)95190-5

Hsieh, S., & Lin, Y.-C. (2016). Stopping ability in younger and older adults: Behavioral and event-related potential. Cognitive, Affective, & Behavioral Neuroscience 2016 17:2, 17(2), 348–363. https://doi.org/10.3758/S13415-016-0483-7

Hsieh, S., & Lin, Y. C. (2017). Strategies for stimulus selective stopping in the elderly. Acta Psychologica, 173, 122–131. https://doi.org/10.1016/J.ACTPSY.2016.12.011

Jana, S., Hannah, R., Muralidharan, V., & Aron, A. R. (2020). Temporal cascade of frontal, motor and muscle processes underlying human action-stopping. ELife, 9. https://doi.org/10.7554/eLife.50371

Kleerekooper, I., van Rooij, S. J. H., van den Wildenberg, W. P. M., de Leeuw, M., Kahn, R. S., & Vink, M. (2016). The effect of aging on fronto-striatal reactive and proactive inhibitory control. NeuroImage, 132, 51–58. https://doi.org/10.1016/j.neuroimage.2016.02.031

Ko, Y.-T., & Miller, J. (2011). Nonselective motor-level changes associated with selective response inhibition: evidence from response force measurements. Psychonomic Bulletin & Review 2011 18:4, 18(4), 813–819. https://doi.org/10.3758/S13423-011-0090-0

Ko, Y. T., & Miller, J. (2013). Signal-related contributions to stopping-interference effects in selective response inhibition. Experimental Brain Research, 228(2), 205–212. https://doi.org/10.1007/s00221-013-3552-y

Lavallee, C. F., Meemken, M. T., Herrmann, C. S., & Huster, R. J. (2014). When holding your horses meets the deer in the headlights: Time-frequency characteristics of global and selective stopping under conditions of proactive and reactive control. Frontiers in Human Neuroscience, 8(DEC), 1–12. https://doi.org/10.3389/fnhum.2014.00994

Lo, S., & Andrews, S. (2015). To transform or not to transform: using generalized linear mixed models to analyse reaction time data. Frontiers in Psychology, 6, 1171. https://doi.org/10.3389/FPSYG.2015.01171/BIBTEX

Logan, G. D., & Cowan, W. B. (1984). On the ability to inhibit thought and action: A theory of an act of control. Psychological Review, 91(3), 295–327. https://doi.org/10.1037/0033-295X.91.3.295

MacDonald, H. J., Coxon, J. P., Stinear, C. M., & Byblow, W. D. (2014). The fall and rise of corticomotor excitability with cancellation and reinitiation of prepared action. Journal of Neurophysiology, 112(11), 2707–2717. https://doi.org/10.1152/jn.00366.2014

MacDonald, Hayley J., McMorland, A. J. C., Stinear, C. M., Coxon, J. P., & Byblow, W. D. (2017). An activation threshold model for response inhibition. PLoS ONE, 12(1), 1–21. https://doi.org/10.1371/journal.pone.0169320

Majid, D. S. A., Cai, W., George, J. S., Verbruggen, F., & Aron, A. R. (2012). Transcranial magnetic stimulation reveals dissociable mechanisms for global versus selective corticomotor suppression underlying the stopping of action. Cerebral Cortex, 22(2), 363–371. https://doi.org/10.1093/cercor/bhr112

Majid, D. S., Cai, W., Corey-Bloom, J., & Aron, A. R. (2013). Proactive selective response suppression is implemented via the basal ganglia. Journal of Neuroscience, 33(33), 13259–13269. https://doi.org/10.1523/JNEUROSCI.5651-12.2013

MathWorks. (2018). MATLAB version 7.10.0. https://au.mathworks.com/products/matlab.html?requestedDomain=

Matzke, D., Love, J., & Heathcote, A. (2017). A Bayesian approach for estimating the probability of trigger failures in the stop-signal paradigm. Behavior Research Methods, 49(1), 267–281. https://doi.org/10.3758/S13428-015-0695-8

Ramautar, J. R., Kok, A., & Ridderinkhof, K. R. (2004). Effects of stop-signal probability in the stop-signal paradigm: the N2/P3 complex further validated. Brain and Cognition, 56(2), 234–252. https://doi.org/10.1016/J.BANDC.2004.07.002

Raud, L., & Huster, R. J. (2017). The Temporal Dynamics of Response Inhibition and their Modulation by Cognitive Control. Brain Topography, 30(4), 486–501. https://doi.org/10.1007/s10548-017-0566-y

Raud, L., Thunberg, C., & Huster, R. J. (2022). Partial response electromyography as a marker of action stopping. ELife, 11. https://doi.org/10.7554/ELIFE.70332

RCoreTeam. (2021). R: A Language and environment for statistical computing. (Version 4.0) [Computer software]. Retrieved from https://Cran.r-Project.Org. (R Packages Retrieved from MRAN Snapshot 2021-04-01).

Schmidt, R., & Berke, J. D. (2017). A Pause-then-Cancel model of stopping: evidence from basal ganglia neurophysiology. Philosophical Transactions of the Royal Society of London. Series B, Biological Sciences, 372(1718). https://doi.org/10.1098/RSTB.2016.0202

Sebastian, A., Konken, A. M., Schaum, M., Lieb, K., Tüscher, O., & Jung, P. (2021). Surprise: Unexpected Action Execution and Unexpected Inhibition Recruit the Same Fronto-Basal-Ganglia Network. Journal of Neuroscience, 41(11), 2447–2456. https://doi.org/10.1523/JNEUROSCI.1681-20.2020

Smittenaar, P., Rutledge, R. B., Zeidman, P., Adams, R. A., Brown, H., Lewis, G., & Dolan, R. J. (2015). Proactive and Reactive Response Inhibition across the Lifespan. PLOS ONE, 10(10), e0140383. https://doi.org/10.1371/JOURNAL.PONE.0140383

Tatz, J. R., Soh, C., & Wessel, J. R. (2021). Towards a two-stage model of action-stopping : Attentional capture explains motor inhibition during early stop-signal processing. BioRxiv. https://doi.org/https://doi.org/10.1101/2021.02.26.433098

The Black Box Toolkit Ltd. [USB response pad]. https://www.blackboxtoolkit.com/urp.html

The jamovi project. (2021). jamovi (Version 2.2). [Computer Software]. Retrieved from https://Www.Jamovi.Org.

van de Laar, M. C., van den Wildenberg, W. P. M., van Boxtel, G. J. M., & van der Molen, M. W. (2011). Lifespan changes in global and selective stopping and performance adjustments. Frontiers in Psychology, 2(DEC), 357. https://doi.org/10.3389/fpsyg.2011.00357

Verbruggen, F., Aron, A. R., Band, G. P. H., Beste, C., Bissett, P. G., Brockett, A. T., Brown, J. W., Chamberlain, S. R., Chambers, C. D., Colonius, H., Colzato, L. S., Corneil, B. D., Coxon, J. P., Dupuis, A., Eagle, D. M., Garavan, H., Greenhouse, I., Heathcote, A., Huster, R. J., … Boehler, C. N. (2019). A consensus guide to capturing the ability to inhibit actions and impulsive behaviors in the stop-signal task. ELife, 8, 1–26. https://doi.org/10.7554/eLife.46323

Verbruggen, F., & Logan, G. D. (2015). Evidence for capacity sharing when stopping. Cognition, 142, 81–95. https://doi.org/10.1016/J.COGNITION.2015.05.014

Wadsley, C. G., Cirillo, J., & Byblow, W. D. (2019). Between-hand coupling during response inhibition. Journal of Neurophysiology, 122(4), 1357–1366. https://doi.org/10.1152/jn.00310.2019

Wadsley, C. G., Cirillo, J., Nieuwenhuys, A., & Byblow, W. D. (2022). Stopping Interference in Response Inhibition: Behavioral and Neural Signatures of Selective Stopping. The Journal of Neuroscience, 42(2), 156–165. https://doi.org/10.1523/jneurosci.0668-21.2021

Wessel, J. R., & Aron, A. R. (2017). On the Globality of Motor Suppression: Unexpected Events and Their Influence on Behavior and Cognition. Neuron, 93(2), 259–280. https://doi.org/10.1016/j.neuron.2016.12.013

