## supplementary materials for "Dissociating Attentional Capture from Action Cancellation in the Stop Signal Task"

**Supplementary Material**

*Preliminary behavioural analysis of stopping interference effects*

To analyse for RT interference effects (i.e., delays in reaction times in successful selective stop, ignore, additional go, and go-ignore trials, relative to the reaction time of the corresponding go response), an initial model was run on data from all conditions including trial type, condition, and age group as factors. Following issues with model convergence (i.e., models still failed to converge when all random effects bar participant intercept were removed) it was determined that running two separate analyses would be the best way to proceed. Two separate GLMMs were run, one including the two SST conditions and another including the two AGT conditions. In the SST conditions, where the stop signal occurred on the right, interference effects were calculated using RTs from the left hand in both bimanual go and right stop trials (Aron & Verbruggen, 2008). Fixed factors included age group (younger, older), condition (SST, SSTignore) and trial type (bimanual go, right stop, right ignore). The ultimate random effects structure included participant intercepts and random slopes for both condition and trial type.

To check for interference effects in the AGTs we used the RT from the response side that was initially cued to go (e.g., the left hand in a left-then-bimanual additional go trial). Fixed factors included age group (younger, older), condition (AGT, AGTignore) and trial type (unimanual go, additional go, go-ignore). Trial type here included only three levels as an initial analysis revealed no significant asymmetry between left- or right-hand responses in either of the AGTs. Thus, the factor of trial type was generalised across hands. That is, left go and right go trials were combined into ‘unimanual go’, left-then-right-go and right-then-left-go trials were combined into ‘additional go’, and left-then-ignore and right-then-ignore trials were combined into ‘go-ignore’. The ultimate random effects structure included participant intercepts and random slopes for both condition and trial type.

Our GLMM checking for interference effects in the SST and SSTignore revealed statistically significant main effects of age ꭓ^2^(1) = 60.24, *p* <0.001, trial type ꭓ^2^(2) = 502.92, *p* <0.001, and condition ꭓ^3^(1) = 5.20, *p* = 0.02. There interaction between condition and age group was not statistically significant ꭓ^2^(1) = 0.08, *p* =0.78, though there were statistically significant interaction effects between trial type and condition ꭓ^2^(1) = 99.50, *p* < 0.001, and between trial type and age group ꭓ^2^(2) = 10.91, *p* = 0.004. There was also a statistically significant three-way interaction between the three variables ꭓ^2^(1) = 9.50, *p* = 0.002. To assess interference effects in each condition, a test of simple main effects of trial type was run, split by condition and age group. In the SST, statistically significant interference effects (the difference between left hand RT in bimanual go and right stop trials) were observed in both the older (*z* = 12.34, *p* <0.001, *d* = 1.10) and younger (*z* = 12.62, *p* < 0.001, *d* = 1.08) cohorts (*see Figure sup1a*). In the SSTignore task, in the older cohort, statistically significant differences between stop and go trials (*z* = 18.17, *p* <0.001, *d* = 1.63) and ignore and go trials (*z* = 11.08, *p* < 0.001, *d* = 0.74) were observed. Similarly, in the younger cohort, statistically significant differences between stop and go trials (*z* = 15.71, *p* <0.001, *d* = 1.31) and ignore and go trials (*z* = 6.53, *p* < 0.001, *d* = 0.46) were observed (*see Figure Sup1b*). Mean RTs can be viewed in Table A2.

The GLMM comparing interference effects in the AGT and AGTignore tasks revealed statistically significant main effects of age ꭓ^2^(1) = 12.26, *p* <0.001, and trial type ꭓ^2^(2) = 68.35, *p* <0.001. The main effect of condition was not statistically significant ꭓ^2^(1) = 0.82, *p* = 0.37. The only interaction to reach statistical significance was between trial type and condition, ꭓ^2^(1) = 14.47, *p* < 0.001. There was no significant interaction between trial type and age ꭓ^2^(2) = 2.40, *p* = 0.30 or between condition and age ꭓ^2^(1) = 0.34, *p* = 0.56. There was also no statistically significant three-way interaction between the three variables ꭓ^2^(1) = 0.10, *p* = 0.75. An exploratory test of simple main effects of trial type was conducted, split by condition and age. In the AGT a small but significant interference effect from additional gos was observed in both the older (*z* = 4.88, *p* < 0.001, *d* = 0.25) and younger (*z* = 3.07, *p* = 0.002, *d* = 0.14) cohorts, with additional go RT slower than the unimanual go RTs. In the AGTignore task, in the older cohort, statistically significant differences between unimanual go and additional go trials (*z* = 6.74, *p* <0.001, *d* = 0.34) and unimanual go and ignore trials (*z* = 3.12, *p* = 0.002, *d* = 0.15) were observed. Likewise, in the younger cohort, statistically significant differences between unimanual go and additional go trials (*z* = 5.27, *p* <0.001, *d* = 0.24) and unimanual go and ignore trials (*z* = 3.90, *p* < 0.001, *d* = 0.17) were observed.

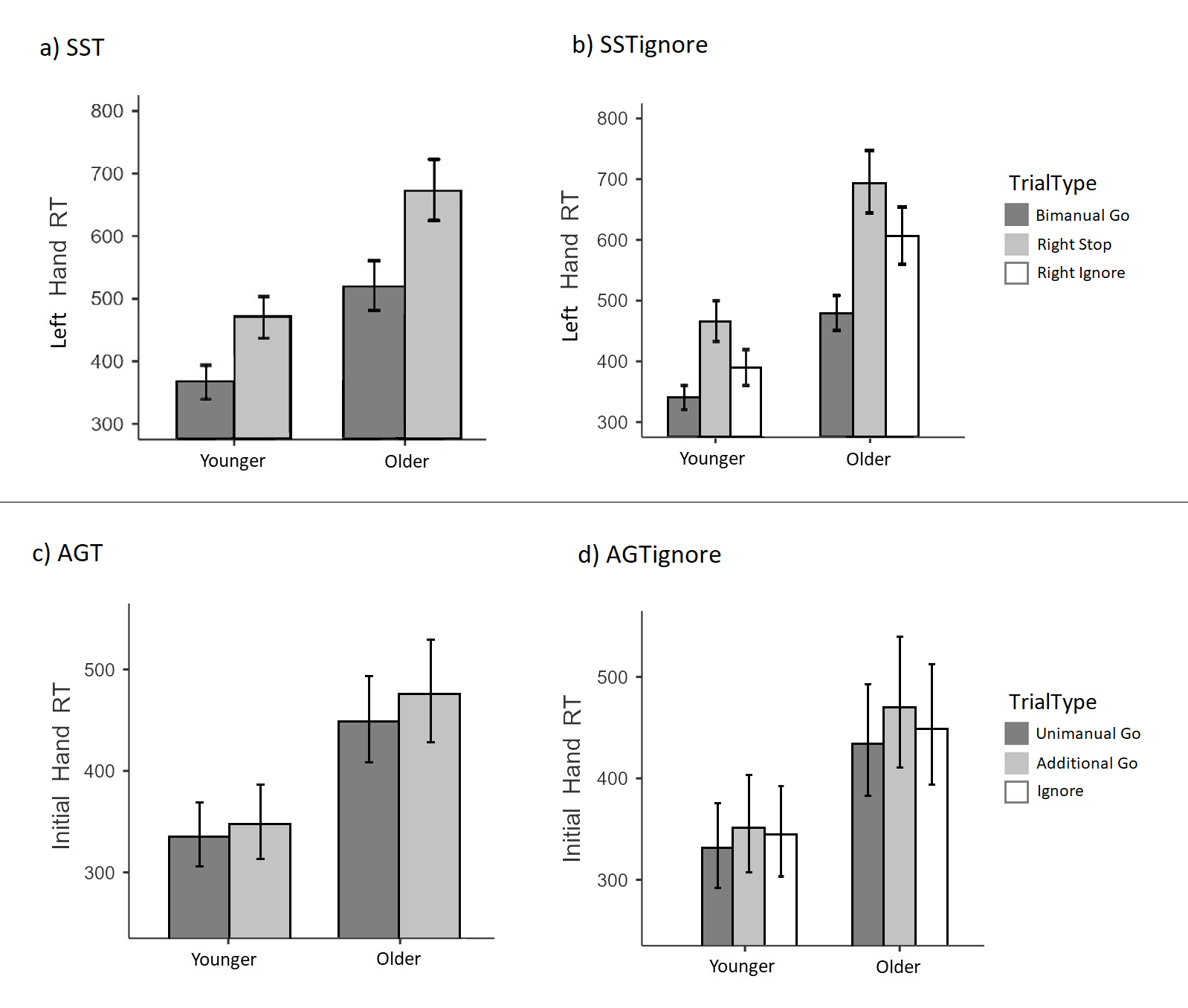

**Figure sup1: mean reaction times (RT; in ms) of the initially responding hand for correct responses in each trial type across each of the four conditions. Simple effects of trial type revealed that, in the SST and SSTignore (panels a and b), right stop and right ignore RTs were significantly longer than bimanual go RTs for both age groups. Furthermore, in the AGT and AGTignore (panels c and d) additional go and ignore RTs were significantly longer than unimanual go RTs, for both age groups. Error bars represent 95%CIs.**

*Preliminary analysis of response amplitude (presence/absence of partial burst not included)*

We used a GLMM approach to analyse the trial-level peak EMG amplitude data. After initial random effects models demonstrated high singularity, two separate models were run (one for the SSTs the other for the AGTs). Preliminary analyses revealed a) no statistically significant difference in peak response amplitude between stop signals in the SST and those in the SSTignore and b) no statistically significant difference between additional go trials in the AGT and AGTignore. As such, the analysis examining SSTs was run on the pooled data from the SST and SST ignore and the analysis run on AGTs was run on the pooled data from the AGT and AGT ignore. For both analyses, trial type and age group were the independent variables and response amplitude was the dependent variable.

The results of the response amplitude analyses are depicted in Figure Sup2. The response amplitude analysis for the two SSTs revealed a statistically significant main effect of age *F*(1,17685) = 22.15, *p* < 0.001. There was also a statistically significant main effect of trial type *F*(2,17685) = 54.68, *p* < 0.001 and a statistically significant interaction effect between age and trial type *F*(2,17685) = 0.89, *p* < 0.001. To investigate the statistically significant interaction an analysis of simple main effects was run on trial type, split by age group. In the older cohort, there was a statistically significant increase in response amplitude in both right stop trials *t*(17685) = 9.83, *p* < 0.001 and ignore trials *t*(17685) = 3.90, *p* < 0.001, relative to bimanual go trials. In the younger cohort, there was a statistically significant increase in response amplitude in right stop trials compared to bimanual go trials *t*(17685) = 5.00, *p* < 0.001; however response amplitude in ignore trials did not significantly differ relative to bimanual go trials *t*(17685) = -0.54, *p* = 0.587.

Our response amplitude analysis for the two AGTs revealed that the main effect of age *F*(1,20527) = 3.77, *p* = 0.052 did not reach statistical significance. There was a statistically significant main effect of trial type *F*(2, 20527) = 5.53, *p* = 0.004 and a statistically significant interaction effect between age and trial type *F*(2, 20527) = 4.56, *p* < 0.011. To investigate the statistically significant interaction an analysis of simple main effects was run on trial type, split by age group. In the older cohort, response amplitude in additional go trials was not significantly different to unimanual go trials *t*(20527) = 0.58, *p* = 0.56; however response amplitude in go-ignore trials *t*(20527) = 4.21, *p* < 0.001 was larger than in unimanual go trials. In the younger cohort, both response amplitude in additional go trials *t*(20527) = 1.50, *p* = 0.13, and go-ignore trials *t*(20527) = -0.30, *p* = 0.76, did not differ from unimanual go trials.

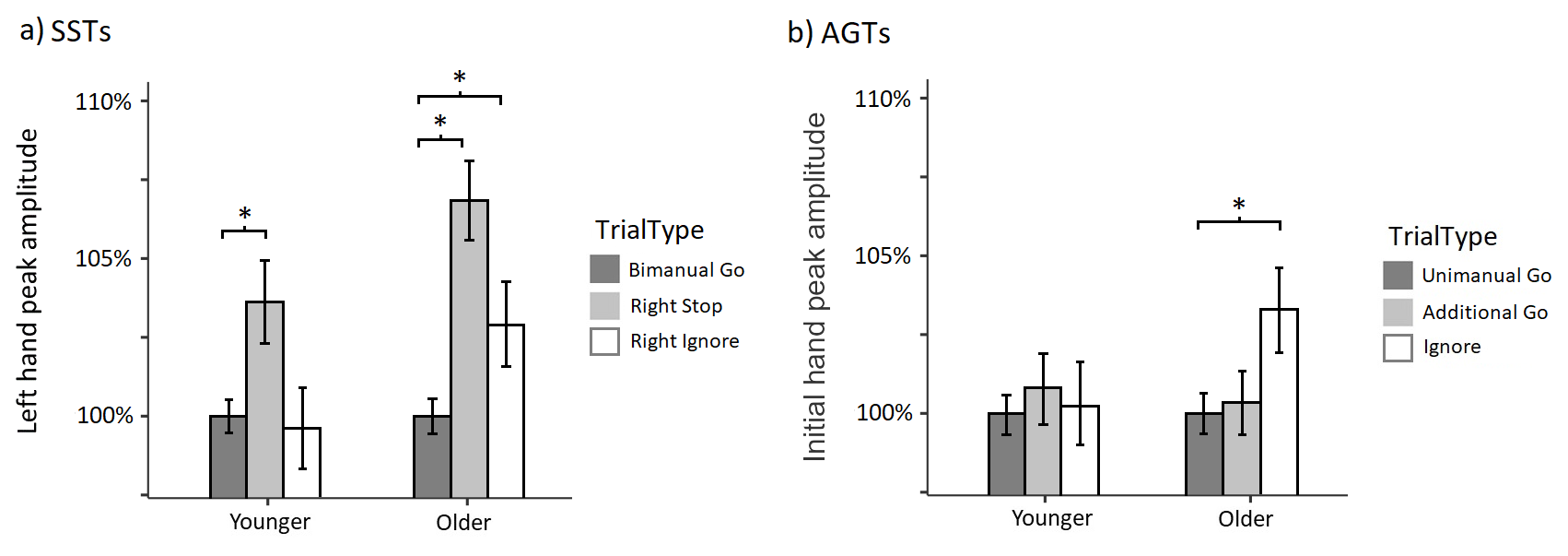

**Figure sup2: Panel a) represents the response amplitude of the non-cancelled (left) hand from the SST and the SSTignore tasks (pooled data). Panel b) represents the response amplitude of the initially cued hand from the AGT and AGTignore tasks (pooled data). Amplitudes are referenced as a percentage of that participant’s average peak EMG levels across all correct go trials in that condition (i.e., amplitude of the unimanual go trials in the AGTs and bimanual go trials in the SSTs are presented as 100% in panel a) and b), respectively). Error bars represent 95%CIs. **p* <0.001.**

Interestingly, we observed an increased response amplitude in ignore trials in the SSTignore, but only in the older cohort. Combined with the observation that the older cohort demonstrated greater behavioural slowing (compared to younger adults) in response to ignore trials (and stop trials), this suggests that stimulus selective inhibition becomes more difficult with advancing age. This may be due to a) a greater latency required for stimulus discrimination processes and/or b) a stopping strategy that prioritises accuracy, leading to a “stop-then-discriminate” strategy (Hsieh & Lin, 2017). Notably, the older cohort also demonstrated increased response amplitude in go-ignore trials in the AGTignore task (Figure sup2). It is conceivable that this finding may also indicate slower stimulus discrimination, whereby a response was initiated following the additional go cue, before discrimination of the fact that the cue should – in fact - be ignored, resulting in the response being cancelled. This could conceivably lead to a raised response threshold and an increased response amplitude in the initially cued hand.

**References (supplementary material)**

Aron, A. R., & Verbruggen, F. (2008). Stop the presses: Dissociating a selective from a global mechanism for stopping: Research article. *Psychological Science*, *19*(11). https://doi.org/10.1111/j.1467-9280.2008.02216.x

Hsieh, S., & Lin, Y. C. (2017). Strategies for stimulus selective stopping in the elderly. *Acta Psychologica*, *173*, 122–131. https://doi.org/10.1016/J.ACTPSY.2016.12.011

**EMG profiles of SST and AGT tasks**

Figures sup3 and sup4 represent the EMG profiles for the SST task for the younger and older cohorts respectively while figures sup5 and sup6 represent the EMG profiles for the AGT task for the younger and older cohorts, respectively. All EMG profiles are synchronised to the onset of the RT generating burst. This allows for clear observation of the partial activations in stop and ignore trials (blue line). Moving the synchronisation reference away from the RT generating burst causes the shape and timing of the EMG profiles to “blur” due to the averaging of profiles whose peaks are not perfectly aligned in time. For this reason, the mean peak EMG in these figures appears to be slightly below 1.

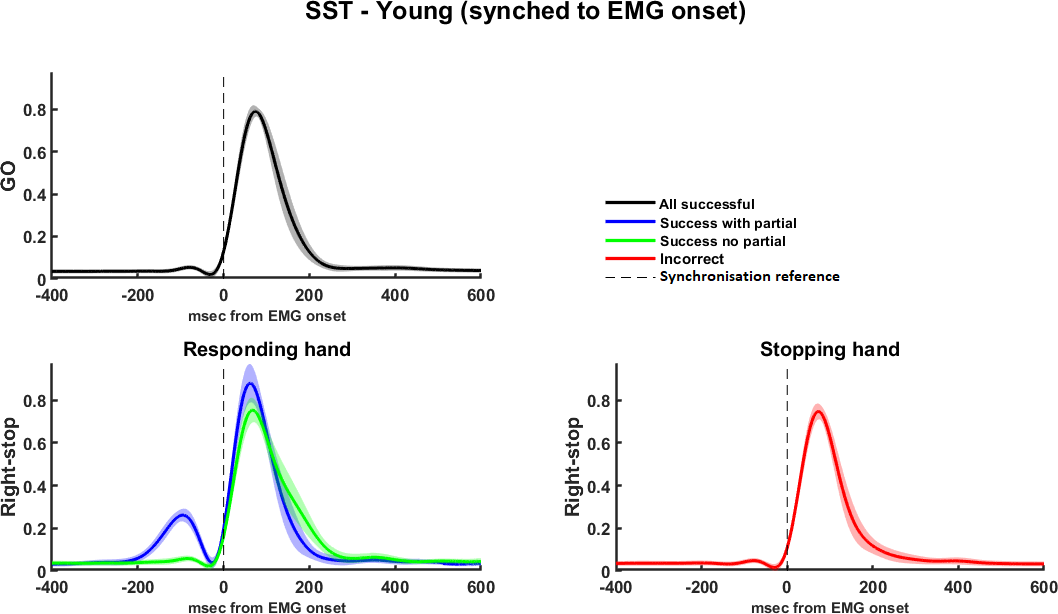

**Figure sup3: EMG profiles for the stop signal task for the younger cohort, split by hand, trial type, and presence/absence of partial burst.** **Trials are synchronised to the onset of the RT generating burst, in order to allow for observation of the presence of partial activations (and the subsequent inhibition). These are most notable on the left hand for right stop and right ignore trials and are represented with the blue line. Shaded areas represent 95%CIs.**

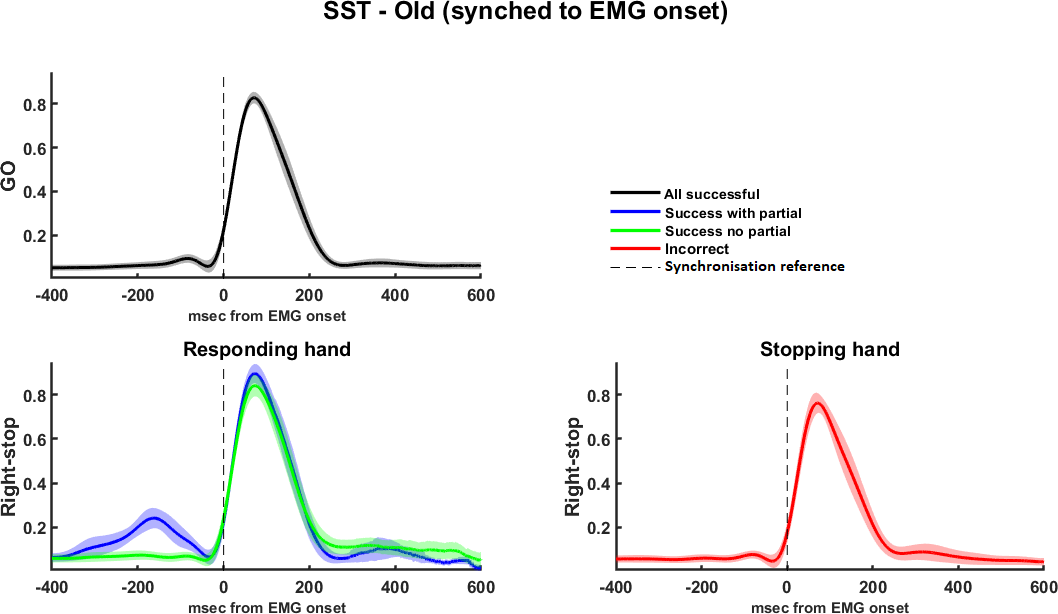

**Figure sup4: EMG profiles for the SST for the older cohort, split by hand, trial type, and presence/absence of partial burst. Trials are synchronised to the onset of the RT generating burst, in order to allow for observation of the presence of partial activations (and the subsequent inhibition). These are most notable on the left hand for right stop and right ignore trials and are represented with the blue line. Shaded areas represent 95%CIs.**

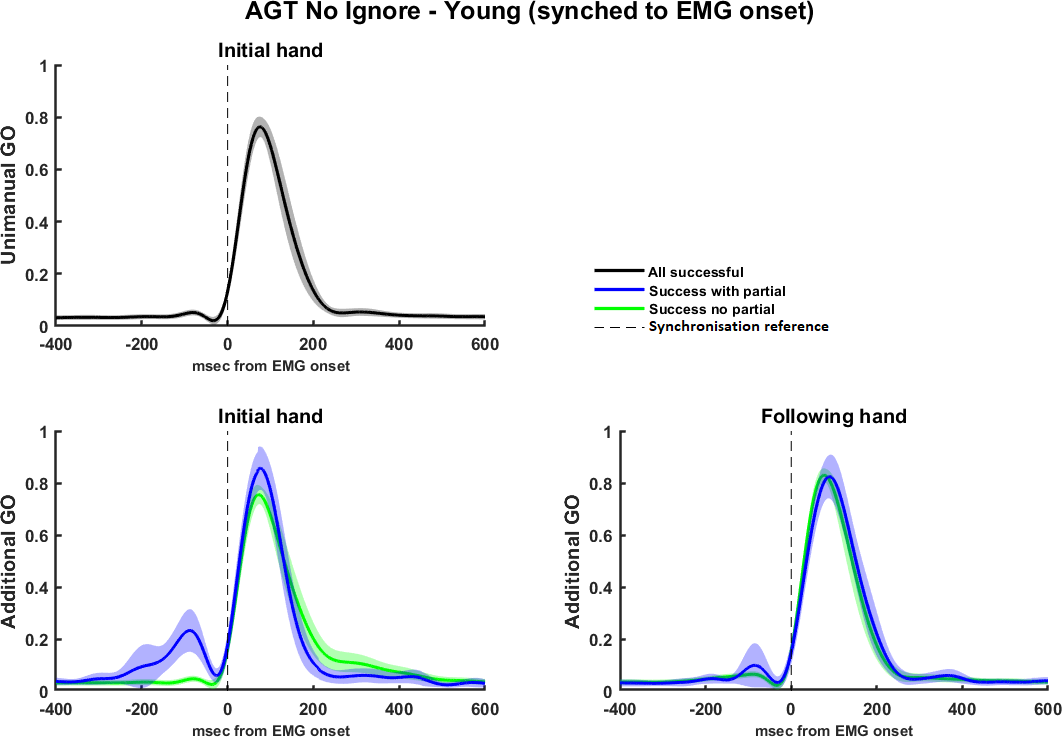

**Figure sup5: EMG profiles for the AGT for the younger cohort, split by hand, trial type, and presence/absence of partial burst. Trials are synchronised to the onset of the RT generating burst, in order to allow for observation of the presence of partial activations (and the subsequent inhibition). These are observable on the left side of left-then-bimanual trials, and the right side of right-then-bimanual trials. Shaded areas represent 95%CIs.**

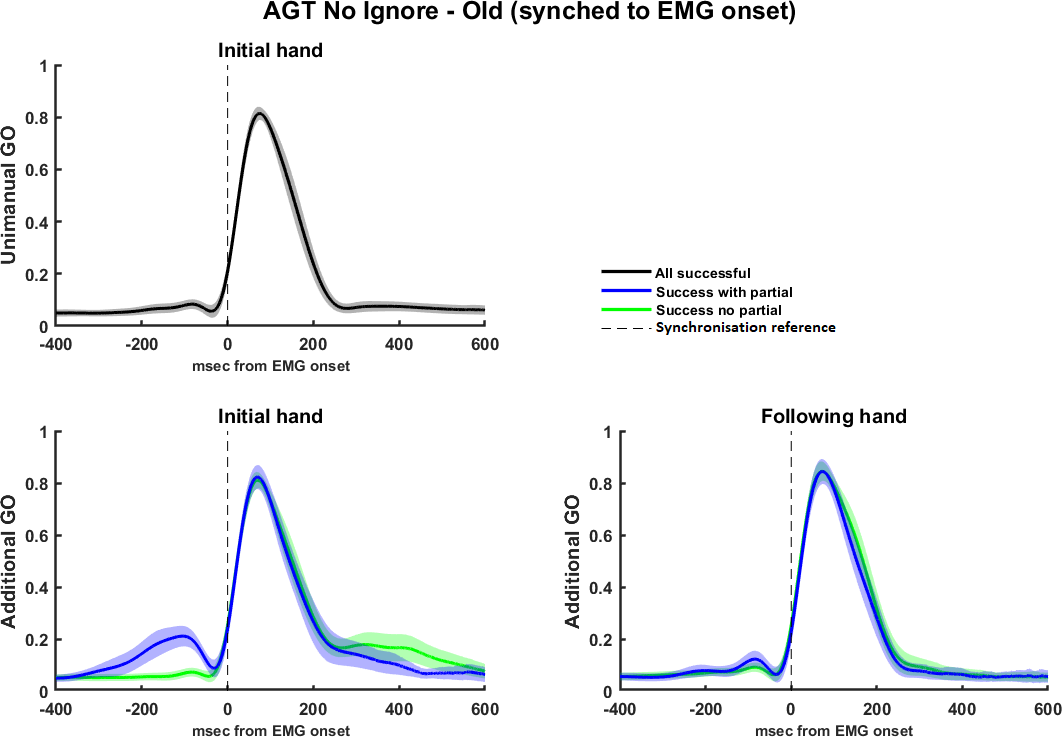

**Figure sup6: EMG profiles for the AGT for the older cohort, split by hand, trial type, and presence/absence of partial burst. Trials are synchronised to the onset of the RT generating burst, in order to allow for observation of the presence of partial activations (and the subsequent inhibition). These are observable on the left side of left-then-bimanual trials, and the right side of right-then-bimanual trials. Shaded areas represent 95%CIs.**
